# Improving the validity of neuroimaging decoding tests of invariant and configural neural representation

**DOI:** 10.1101/2020.02.27.967505

**Authors:** Fabian A. Soto, Sanjay Narasiwodeyar

## Abstract

Many research questions in sensory neuroscience involve determining whether the neural representation of a stimulus property is invariant or specific to a particular stimulus context (e.g., Is object representation invariant to translation? Is the representation of a face feature specific to the context of other face features?). Between these two extremes, representations may also be context-tolerant or context-sensitive. Most neuroimaging studies have used operational tests in which a target property is inferred from a significant test against the null hypothesis of the opposite property. For example, the popular cross-classification test concludes that representations are invariant or tolerant when the null hypothesis of specificity is rejected. A recently developed neurocomputational theory provides two insights regarding such tests. First, tests against the null of context-specificity, and for the alternative of context-invariance, are prone to false positives due to the way in which the underlying neural representations are transformed into indirect measurements in neuroimaging studies. Second, jointly performing tests against the nulls of invariance and specificity allows one to reach more precise and valid conclusions about the underlying representations. Here, we provide empirical and computational evidence supporting both of these theoretical insights. In our empirical study, we use encoding of orientation and spatial position in primary visual cortex as a case study, as previous research has established that these properties are encoded in a context-sensitive way. Using fMRI decoding, we show that the cross-classification test produces false-positive conclusions of invariance, but that more valid conclusions can be reached by jointly performing tests against the null of invariance. The results of two simulations further support both of these conclusions. We conclude that more valid inferences about invariance or specificity of neural representations can be reached by jointly testing against both hypotheses, and using neurocomputational theory to guide the interpretation of results.

**Author Summary:** Many research questions in sensory neuroscience involve determining whether the representation of a stimulus property is invariant or specific to a change in stimulus context (e.g., translation-invariant object representation; configural representation of face features). Between these two extremes, representations may also be context-tolerant or context-sensitive. Most neuroimaging research has studied invariance using operational tests, among which the most widely used in recent years is cross-classification. We provide evidence from a functional MRI study, simulations, and theoretical results supporting two insights regarding such tests: (1) tests that seek to provide evidence for invariance (like cross-classification) have an inflated false positive rate, but (2) using complementary tests that seek evidence for context-specificity leads to more valid conclusions.

## Introduction

A common question in sensory and cognitive neuroscience is to what extent the neural representation of a stimulus property changes as a function of changes in other aspects of stimulation–that is, the context in which it is presented–. As shown in Figure 1, one possibility is that the neural representation of the target property is invariant to changes in context. In that case, the neural activity representing the target property does not change at all with changes in context. Another possibility is that the neural representation of the target property is context-specific. In that case, the neural activity representing the target property completely changes with a change in context. Another way to describe context-specificity is by saying that the target property and its context are represented configurally; that is, as a configuration separate from its components. As shown in Figure 1, these two cases of complete invariance and specificity should be seen as extremes in a continuum. In this continuum, representations that are closer to invariance could be characterized as “tolerant” to changes in context, whereas representations that are closer to specificity could be characterized as “sensitive” to changes in context.

**Figure 1:**
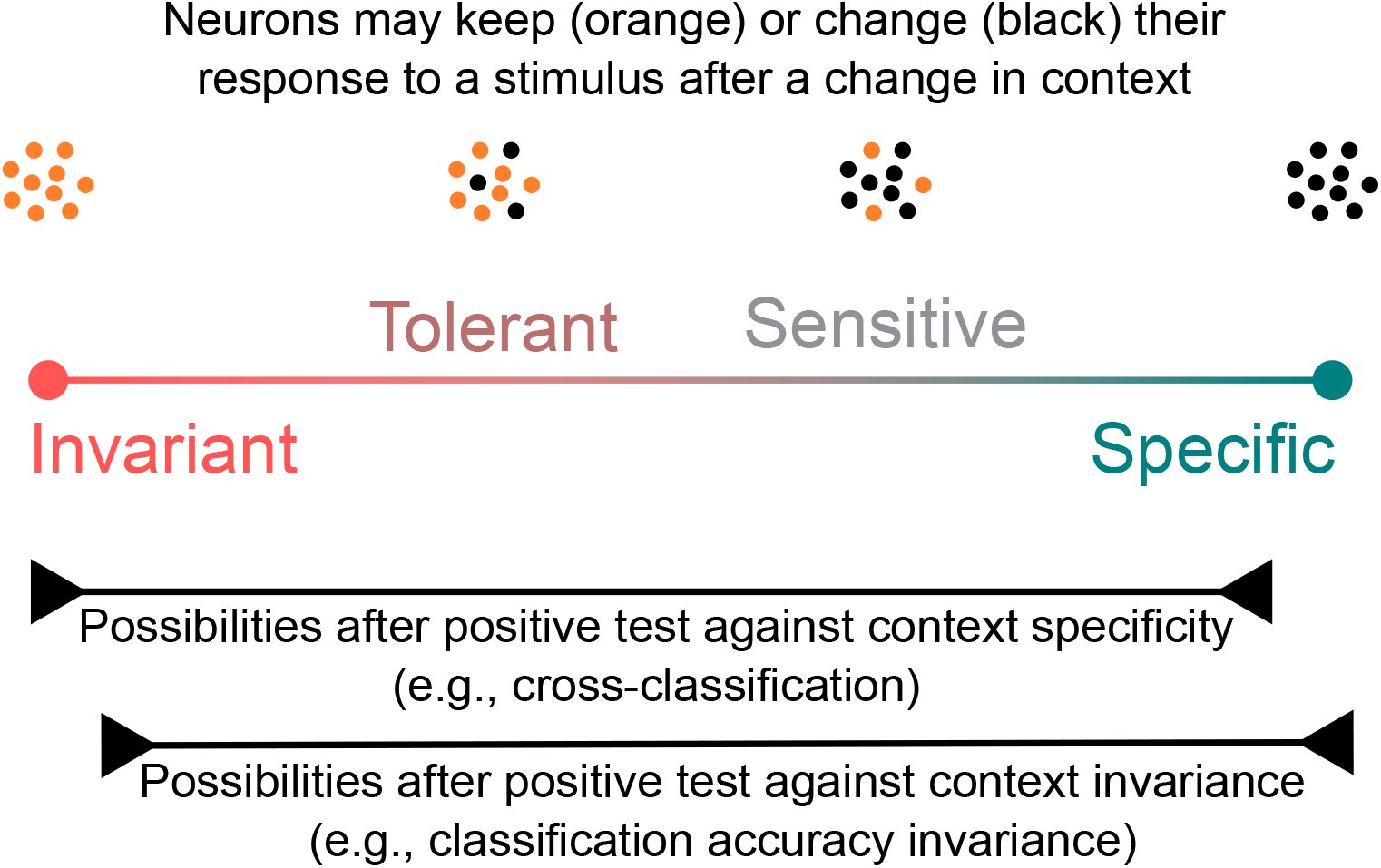
Encoding of a target feature by a group of neurons can have varying degrees of change as a function of a change in context. With a change in context, context-invariant representations do not change at all, whereas context-specific representations change completely. There is a continuum between both extremes, which includes the more likely cases of context-tolerance and context-sensitivity. A significant test against the null hypothesis of context specificity, such as that provided by the cross-classification test, is usually interpreted as favoring invariance although the representation can be anywhere from sensitive to invariant. Similarly, a significant test against the null of context invariance can be obtained from representations ranging from tolerant to specific. We propose that jointly performing both types of test considerably increases the validity of our conclusions regarding the underlying representations.

Most human neuroimaging research has studied invariance and specificity using operational tests that provide evidence against the null hypotheses represented by the two extremes in Figure 1. This choice is paradoxical, as most neuroscientists are more interested in determining to what extent a representation is closer to one of the extremes in the continuum, being classified as either context-tolerant or context-sensitive.

For example, probably the most widely used test in this area is cross-classification (or cross-decoding; 1, 2, 3; we have also called this test classification accuracy generalization: [4]), illustrated in Figure 2. The first step in cross-classification is to train a classifier to decode a particular stimulus feature, such as whether a presented face is male or female, from patterns of fMRI activity observed across voxels. The second step is to test the trained classifier with new patterns of fMRI activity, this time obtained from presentation of the same stimuli, but changed in an irrelevant property, such as head orientation. Using our nomenclature, in this example the target stimulus property is face sex, and the context is face orientation. If accuracy with the test data is higher than chance, then researchers usually conclude that the neural representation of the target feature has a certain level of tolerance to changes in context (usually described as invariance), within the area from which the fMRI activity was obtained.

**Figure 2:**
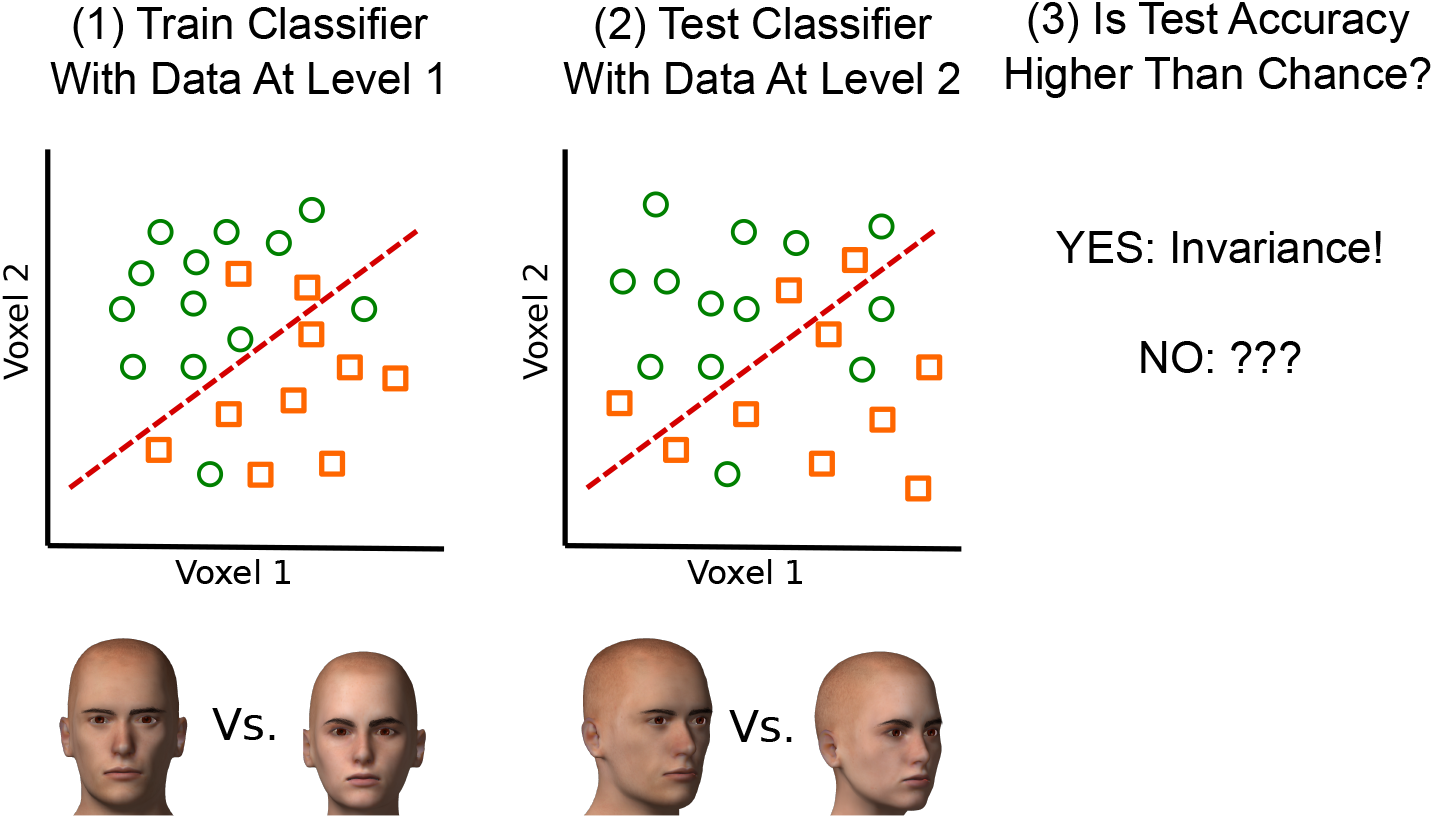
Cross-classification test. (1) A classifier is trained to discriminate between levels of a target dimension (facial gender) in a given level of the context dimension (viewpoint). The axes represent activity in two exemplary voxels in response to stimuli. Each data point represents a single trial. (2) The same classifier is then used to classify data from a second level of the context dimension (different viewpoint). (3) If classification accuracy is above chance levels across viewpoints, then invariance is said to hold according to this test.

The cross-classification test has been used to provide evidence for tolerant encoding of face identity across viewpoint [5], object category and viewpoint across spatial position [6], object category across shape [and vice-versa; 7], motor actions across modalities [8], place of speech articulation features across manner of articulation [9], object category [10] or face identity [11] across stimulus modality, word semantic category across stimulus modality [12], learned category labels across categorization tasks [13], and semantic word representation across languages [14], among others [for a review, see 3].

Cross classification is a test against the null hypothesis of no generalization of decoding accuracy from one context to another, a condition that would be met under context-specific encoding of the target property. As shown in Figure 1, evidence against the extreme of context-specificity means that the representation can fall anywhere in the continuum except the right extreme. Invariance and tolerance are only some of the possibilities, as representations may also be context-sensitive.

An example of a test that provides evidence against the null hypothesis of invariance is the *classification accuracy invariance* test [4]. This test involves the same steps described for cross-classification in Figure 2, but during the test phase the classifier is presented with data obtained at both the training and the testing context levels (i.e., levels 1 and 2 in Figure 2). The null hypothesis is that decoding accuracy is equivalent across all levels of context. When accuracy drops significantly from training to testing context, one can conclude that the underlying representation of the decoded property is not invariant to context. While the test was independently developed from theory [4], we are aware of at least one prior study using a version of this test to obtain evidence of position-dependent encoding of object category information in lateral occipital cortex, and of position-dependent encoding of face viewpoint information in right fusiform face area [6].

Again, evidence against the extreme of context-invariance means that the representation can fall anywhere else in the continuum shown in Figure 1. Context specificity is only one of the possibilities, as representations may also be context-sensitive or context-tolerant.

In a previous theoretical paper [4], we explored to what extent the context tolerance or specificity of neural representations could be measured using a variety of neuroimaging analyses, with a focus on decoding tests like cross-classification and classification accuracy invariance. Because neuroimaging involves only indirect measures of neural activity, it cannot be used to get precise indicators of where a neural representation falls within the continuum shown in Figure 1. In general, the process by which neural representations are transformed from the neural space into a space of measurements (e.g., voxel activities) will distort the representations in such a way that makes such precise indicators impossible. However, the results of neuroimaging decoding tests like those just described do allow to make some inferences about the underlying neural representations. Besides clarifying what different tests measure (i.e., cross-classification provides evidence *against* context-specificity, rather than evidence *for* invariance), this theoretical work provides two important insights that have consequences for the way in which neuroimaging researchers should apply and interpret the results of decoding tests.

The first theoretical insight is that jointly performing tests against the nulls of invariance and specificity allows one to reach more precise and valid conclusions about the underlying representations. When both types of tests are carried out, one can use Table 1 to reach valid conclusions about properties of the underlying neural code. For example, one may use the cross-classification test to obtain evidence against context-specificity, but usually researchers who use this test are interested in reaching a conclusion favoring invariance or tolerance [e.g., 5, 3]. For that, information from a test against invariance would be very useful. If a test against invariance is not significant, one can make a stronger case for tolerant representations. Because sample size and measurement noise are equivalent in this test and the significant cross-classification test, the best interpretation is that the underlying representation is likely to be farther away from specificity than from invariance, being tolerant/invariant rather than sensitive. On the other hand, if the test offers evidence against invariance, then the underlying representations could be anywhere in the continuum shown in Figure 1, except at the two extremes, and it would be premature to make a conclusion of tolerance in the underlying representations, as they are equally likely to be context-sensitive. Because tests against invariance have been rarely used in the literature, one goal of the current study is to provide evidence of the validity of such tests.

**Table 1:** Lookup table summarizing how joint tests against specificity and invariance should be interpreted. Note that significance of the popular cross-classification test does not guarantee a conclusion for tolerance or invariance. Only when such a test is accompanied by a nonsignificant test against invariance one can reach a positive conclusion.

|  |  | Test against specificity (e.g., cross-classification) |  |
| --- | --- | --- | --- |
|  |  | Not significant | Significant |
| Test against invariance (e.g., decoding separability) | Not significant | Inconclusive results | Tolerance or invariance likely |
|  | Significant | Specificity or sensitivity likely | Inconclusive results |

The second theoretical insight is that there is an important asymmetry regarding the validity of tests of invariance and context-specificity. If the underlying neural representation is truly invariant, then a signal showing evidence against invariance will never be found from neuroimaging decoding tests. In this case, any finding of lack of invariance would result from measurement noise, and the probability of such finding would be equal to the false positive (type I) error rate of the statistical test, usually *α* = .05. On the other hand, if the underlying representation is truly context-specific, it is still possible to find a signal at the level of voxels showing evidence against context-sensitivity. In this case, such a signal will add to the probability of false positives, which would be higher than *α*.

The reason lies in the contribution of the measurement model, which summarizes how representations are transformed from the space of neural representations into the space of measured variables. Figure 3 depicts a schematic example, where representations of the target stimuli in one context (e.g., faces with front orientation) are shown as red circles, and representations in a second context (e.g., faces with sideways orientation) are shown as green crosses. In panel (a), the original neural representations are fully context-invariant, meaning that the representation of a stimulus in either context is in the exact same point in neural space. Regardless of what transformation is induced by the measurement model, such representations will remain invariant in the measurement space, as the transformation will have the same effect on two identical representations (i.e., overlapping crosses and circles in Figure 3). In panel (b), the original representations are fully context-specific, meaning that the stimulus representations occupy completely different regions of space depending on context. In this case, there are transformations that would reduce differences in the representation of stimuli across contexts, making the representations less context-specific. In sum, the transformation from neural space to measurement space (i.e., the measurement model) cannot make a completely invariant representation appear as if it was sensitive to context, but it can make a completely context-specific representation appear as if it was tolerant to changes in context.

**Figure 3:**
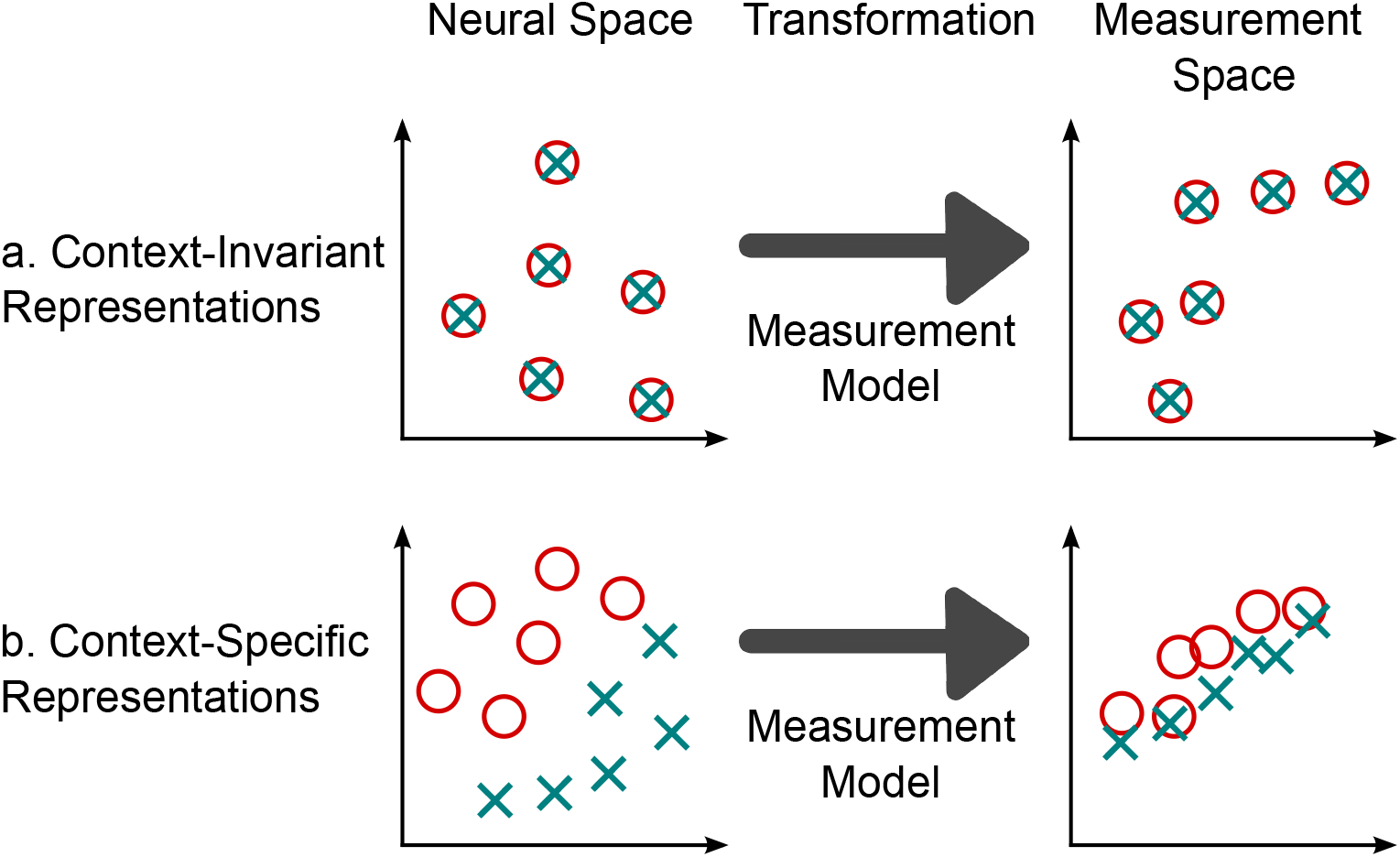
A measurement model represents how representations are transformed from the space of neural activities to the space of voxel measurements. The outcome of this transformation has an asymmetrical effect on context-invariant and context-specific representations. Context-invariant representations (a) cannot be transformed in such a way to decrease their invariance and increase their specificity. On the other hand, context-specific representations (b) can be transformed to increase their invariance and decrease their specificity. The result is that there is an inflated risk to find false positive invariance in neuroimaging studies such as those using the cross-classification test.

This can happen in a number of ways, but the simplest example is one in which encoding of the target property is spatially smooth across voxels, while changes in context produce changes in the spatial distribution of activity that are fine-grained [15]. Take the example shown in Figure 4. Each column in the figure represents a different voxel, which itself contains a large number of neurons (or neural populations), represented by small circles, with selectivity for some target stimulus property. In this simplified example, the neurons can show preference for one of two values of the target property, represented by the colors red and yellow. Neurons can be inactive in a particular context, which is represented by the color gray. Different voxels have different proportions of the two types of neurons, so that despite of the spatial pooling of activity produced at each voxel, there is a distinctive pattern of activity produced across voxels by each stimulus property. This is a spatially smooth coding scheme.

**Figure 4:**
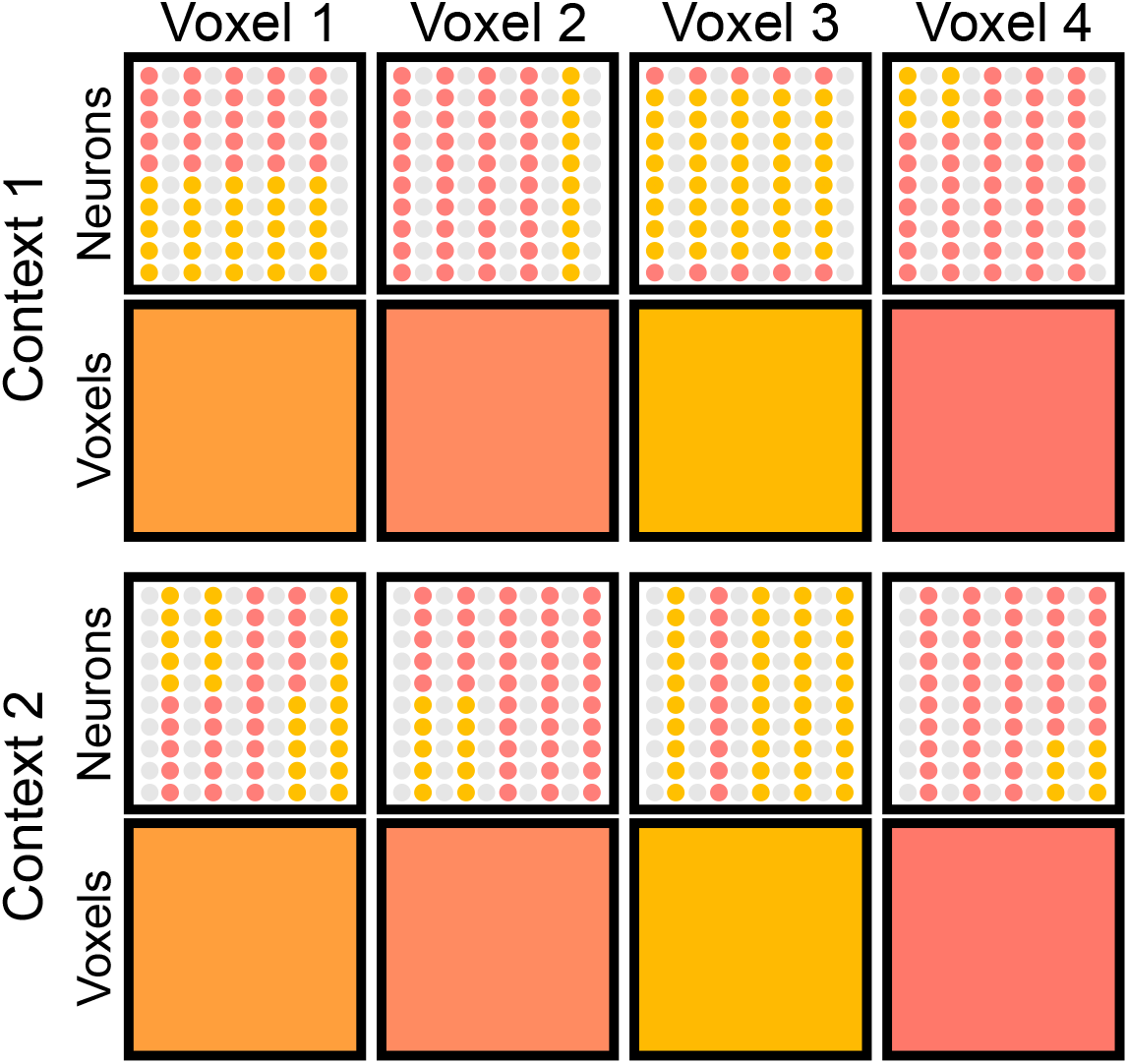
Example highlighting the differences between spatially smooth versus fine-grained encoding schemes, and a particular combination of the two schemes that produces false-positives in a voxelwise analysis. Each column in the figure represents a different voxel containing a large number of neurons represented by small circles, each with selectivity for one of two values of the target property, represented by the colors red and yellow. Different proportions of the two types of neurons are present across different voxels, so that despite of the spatial pooling of activity produced at each voxel, there is a distinctive pattern of activity produced across voxels by each stimulus property. This is a spatially smooth coding scheme. On the other hand, within a voxel widely different spatial distributions of activity may produce the same value of global activity at the voxel level, in a fine-grained coding scheme. Note how the multivoxel pattern of activity is the same for both levels of the context dimension, even though completely different populations of neurons encode each level. The combination of spatially smooth encoding of the target dimension and fine-grained encoding of the context dimension would produce false-positive invariance in a voxelwise analysis.

On the other hand, note how within a voxel widely different spatial distributions of activity may produce the same value of global activity at the voxel level. For example, the same aggregate activity is obtained for voxel 1 in context 1 (top) and context 2 (bottom), despite the fact that the fine-grained distribution of activities is widely different. The same is true for all other voxels. Thus, within each voxel one can see a fine-grained coding scheme that distinguishes between contexts.

More importantly for the issue of neural invariance, in Figure 4 the neurons encoding the target dimension in the first context (uneven columns of neurons) are completely different to those encoding the target dimension in the second context (even columns of neurons). However, the spatial distribution of neurons specific to each value of the context dimension is spatially homogeneous, with about the same number of neurons of each kind in the voxel regardless of context.

The result of a spatially smooth encoding of the target dimension across voxels, together with a fine-grained spatial distribution of neurons specific to each value of the context dimension, produce as a result a case in which neural encoding of the target dimension is context-specific, but appears as perfectly invariant at the level of voxel activities.

A good example of this type of encoding in the brain is encoding of spatial position and orientation in V1. Encoding of spatial position is spatially smooth in V1, with the scale of retinotopic maps being similar to the voxel sizes typically used in neuroimaging, whereas encoding of orientation is much more spatially fine-grained [see 16, 17]. Indeed, orientation maps are so fine-grained compared to the spatial scale of voxels in typical experiments that researchers have debated for years how it is that we are able to decode orientation from V1 using fMRI in the first place [e.g., 18, 19, 20].

This example shows that the kind of encoding scheme exemplified by Figure 4 can be found in the brain. Because of the influence of the measurement model depicted in Figure 3b, the need to jointly perform and interpret tests of invariance and specificity is even greater for researchers who aim to find evidence for tolerant/invariant representations. If a false positive is found in a test of context-specificity (e.g., cross-classification) due to issues in the measurement model, it is unlikely that a test of invariance (e.g., classification accuracy invariance) will also be significant. That is, while the cross-classification test has an inherent tendency to produce more false positives than expected (i.e., *> α*), this issue can be partially controlled by interpreting the results of that test together with results of tests against the null of invariance.

Here, we show that the two theoretical insights described above have important consequences for neuroimaging research, through empirical evidence coming from an fMRI decoding study, and computational evidence coming from simulation work. In the empirical study, we perform decoding of orientation and spatial position from fMRI activity patterns recorded in V1, a case in which properties of the underlying neural code are known. The cross-classification test provides strong evidence for the incorrect conclusion that, in V1, encoding of spatial position is tolerant/invariant to changes in orientation, as well as some evidence for the incorrect conclusion that orientation is tolerant/invariant to changes in spatial position. We find that the use of theoretically-derived tests of invariance can lead to more valid conclusions regarding the underlying code. The results of two simulations further support all of these conclusions. Our results highlight the validity and value of using tests of invariance together with tests of context-specificity (e.g., cross-classification) when attempting to draw inferences about neural representations from neuroimaging decoding studies.

## Results

### Experimental Results

The goal of our study was to validate two insights provided by neurocomputational theory [4]. First, that tests aimed at testing against the null hypothesis of context-specificity, and to provide evidence for the alternative hypothesis of context-invariance, may be prone to false positives due to the way in which the underlying neural representations are transformed into measurements, as shown in Figures 3 and 4. Second, that jointly performing tests against the nulls of invariance and specificity allows one to reach more precise and valid conclusions about the underlying representations.

To test these two hypotheses, we applied decoding tests of invariance and specificity to the study of orientation and spatial position in V1. Previous research has established that these properties are not encoded in an invariant way but, as explained in the introduction, the spatial scale of orientation and spatial position maps in V1 is likely to lead to the incorrect conclusion of invariance if tests of specificity, such as cross-classification, are applied on their own.

Participants were presented with the stimuli in Figure 13 while they performed a task involving a stimulus presented at the center of the screen. Functional MRI data was acquired at the same time, with separate runs providing data for training and testing of a support vector machine (SM) classifier. Training runs were composed of stimuli presented only in spatial positions top-right and bottom-left (highlighted through red and blue boxes in Figure 13). Testing runs included all sixteen stimulus combinations. We trained a linear SVM classifier to decode a target dimension (e.g., spatial position) while holding the context dimension (e.g., grating orientation) constant. We then tested the classifier with data obtained at the trained value of the context dimension (e.g., 0° orientation) as well as new values of the context dimension (e.g., 45°, 90°, and 135° orientation). The classifier provided decision variables and accuracy estimates used to perform a test of specificity (cross-classification) and two tests of invariance (classification accuracy invariance, decoding separability) presented below (for more details, see *Materials and Methods*).

### The Cross-Classification Test Produces False Positives

The popular cross-classification test ([e.g., 1, 2, 3], see Figure 2) was specifically designed to provide evidence in favor of invariant encoding in a given brain region, by testing against the null of context-specific encoding (see Figure 1). As indicated earlier (see Figures 3 and 4), theoretical considerations suggest that such a test would be biased to generate false positives, leading to evidence against context-specificity that is due to distortions in neural representations imposed by the measurement model, rather than to actual properties of neural encoding (see Figure 3b). We performed a set of analyses using the cross-classification test to validate our theoretical prediction that this method should produce findings of false-positive invariance; that is, cases in which invariance is concluded even though our knowledge of the underlying neural code indicates that such invariance does not exist. The cross-classification test was conducted by assessing whether a linear decoder trained to classify the target dimension at one level of the context dimension, could perform the same classification above chance across non-trained levels of the context dimension. A positive result in the cross-classification test is usually taken as evidence for the existence of invariant representations in the area of interest [2, 3].

We conducted two separate analyses using the cross-classification test in which we switched the identities of the target and context dimensions. In the first analysis, spatial position was treated as the target dimension to be decoded, while orientation remained as the context dimension. To obtain decoded stimulus values for spatial position, we used deconvolved single-trial estimates of activity in V1 voxels as input to the SVM linear decoder. We trained the decoder to classify trials based on spatial position labels (top-right vs bottom-left, see boxed stimuli in Figure 13) and holding constant the level of grating orientation (context dimension; for example, 0°) using leave-one-run-out cross-validation, and tested it with independent data sets across all levels of grating orientation (0°, 45°, 90°, and 135°). To test for cross-classification invariance, we performed a binomial test on the accuracy estimates from the testing data set, corrected for multiple comparisons using the Holm-Sidak method. (for more details, see *fMRI Decoding Tests*) If the accuracy score was significantly above chance, then the cross-classification test concludes that spatial position is encoded invariantly from orientation in V1, a conclusion known to be false.

For each participant, we repeated the analysis four times, once for each level of grating orientation that was held fixed in the classifier’s training data. Based on theoretical considerations [4], we predicted that the cross-classification test would generate consistent false positives in the case where spatial position was used as the relevant dimension to be decoded. Since spatial position is encoded in a spatially smooth manner in V1, we expected strong performance of the classifier across all levels of orientation. In other words, we expected the accuracy scores of the classifier to remain above chance across different levels of the context dimension.

Figure 5 shows accuracy estimates from such a decoding procedure for all five subjects. The SVM linear decoder achieves extremely high levels of classification accuracy in test sets across all 5 subjects. As predicted, the test incorrectly finds evidence for invariance of spatial position from orientation in all participants and all tests (all *p*<.001; for details see Table 1 in the Supplementary Material). This result is unsurprising, in the sense that one would intuitively expect it given the properties of encoding in V1, where information about the spatial position of stimuli is spatially smooth, distributed at around the same scale as our voxel size, but information about orientation is fine-grained, distributed at a smaller scale than our voxel size. The important point, however, is that in most applications of the cross-classification test researchers do not know much about encoding in the area under study, and they could easily conclude in favor of invariance when the underlying code does not show such property.

**Figure 5:**
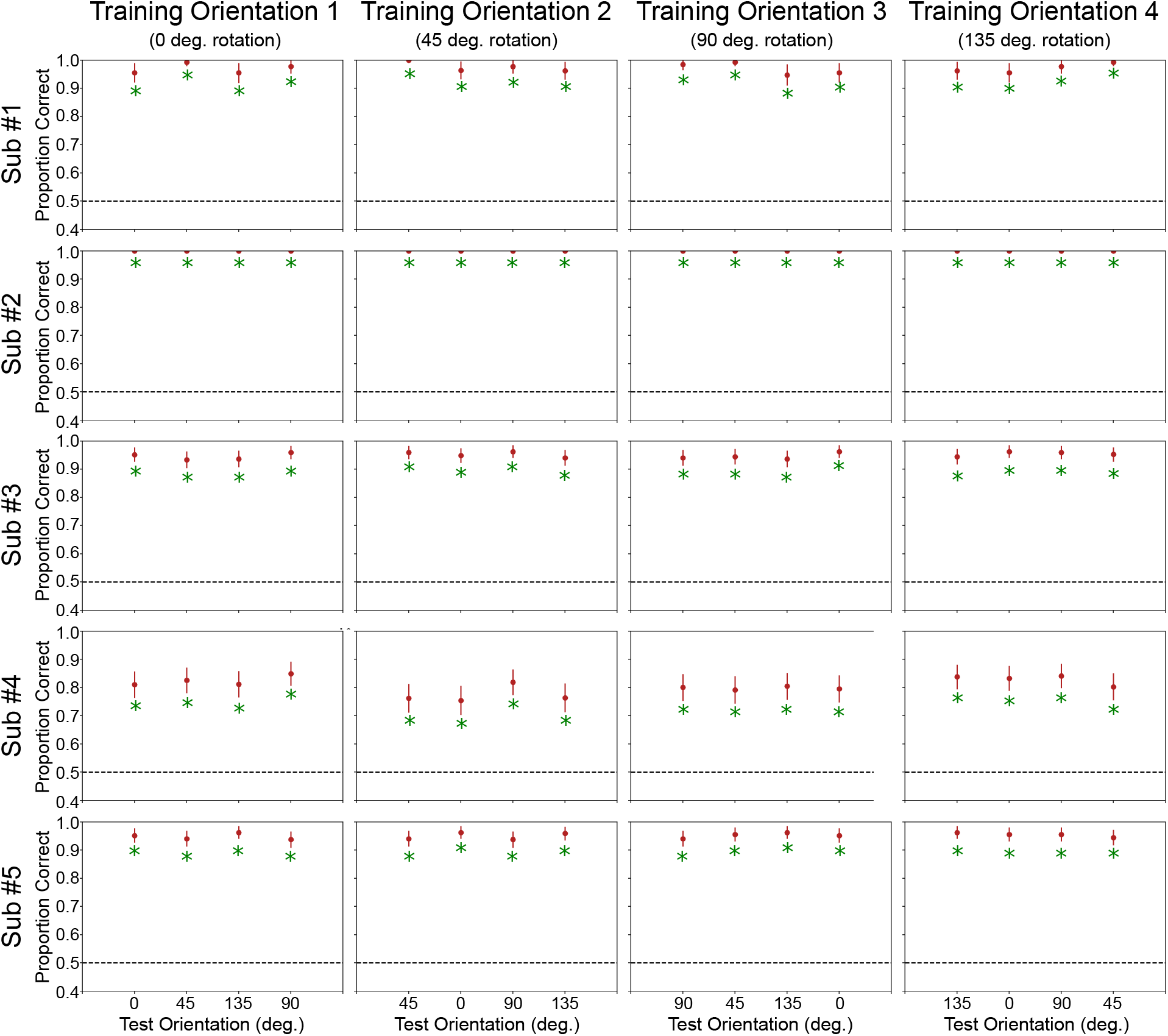
Classification accuracy results with test data for the decoding of spatial position. Each row represents a complete analysis for a single subject. Columns represent different levels of the context dimension (orientation) held fixed during training. For example, the leftmost cell for subject 1 displays accuracy results from a decoder that was trained to discriminate spatial positions while the oriented grating was held constant at 0°. The same decoder was then tested with independent testing data in which the oriented grating was held at all orientations (0°, 45°, 90°, and 135°). A green asterisk represents accuracy values significantly above chance (dotted line), leading to the conclusion of invariance from the cross-classification test. As expected, the cross classification test consistently generates false positives and yields a conclusion that spatial position is encoded invariantly from orientation in V1.

We performed a second analysis in which orientation was treated as the target dimension to be decoded, while spatial position was treated as the context dimension. We trained the decoder to classify trials based on grating orientation (0°, 45°, 90°, and 135°, see boxed stimuli in Figure 13) and holding constant the position of the spatial window (context dimension; for example, top-right or 20° in Figure 13) using leave-one-run-out cross-validation, and tested it with independent data sets across all levels of spatial position (top-right, bottom-right, bottom-left, and top-left; or 20°, 80°, 200°, and 260° in Figure 13). All other procedures remained the same as in the first analysis. Figure (6) shows decoding accuracy results for the orientation analysis. The SVM classifier was able to successfully decode orientation information at the original training position in all subjects, but for subjects 1 and 4 this was restricted to a single training window (200° window). In contrast to spatial position classification, the classifier’s accuracy scores drop significantly in untrained testing windows.

The classifier accuracy at the training window provides a ceiling of performance for the cross-classification accuracy (see 2, 3). That is, we are not interested in the analyses with non-significant accuracies at the training window (sub#1 and sub#4 at training window 1; see Figure 6), as in those cases we would not expect a significant cross-classification accuracy. Out of the eight analyses showing significant accuracy at the training window, two generated significant cross-classification results, which would lead to an invalid conclusion of invariance. This number was higher than the 5% expected false positive rate for these tests, but a binomial test did not reach significance with *p* =.051, probably due to the low power of a test involving only eight analyses.

**Figure 6:**
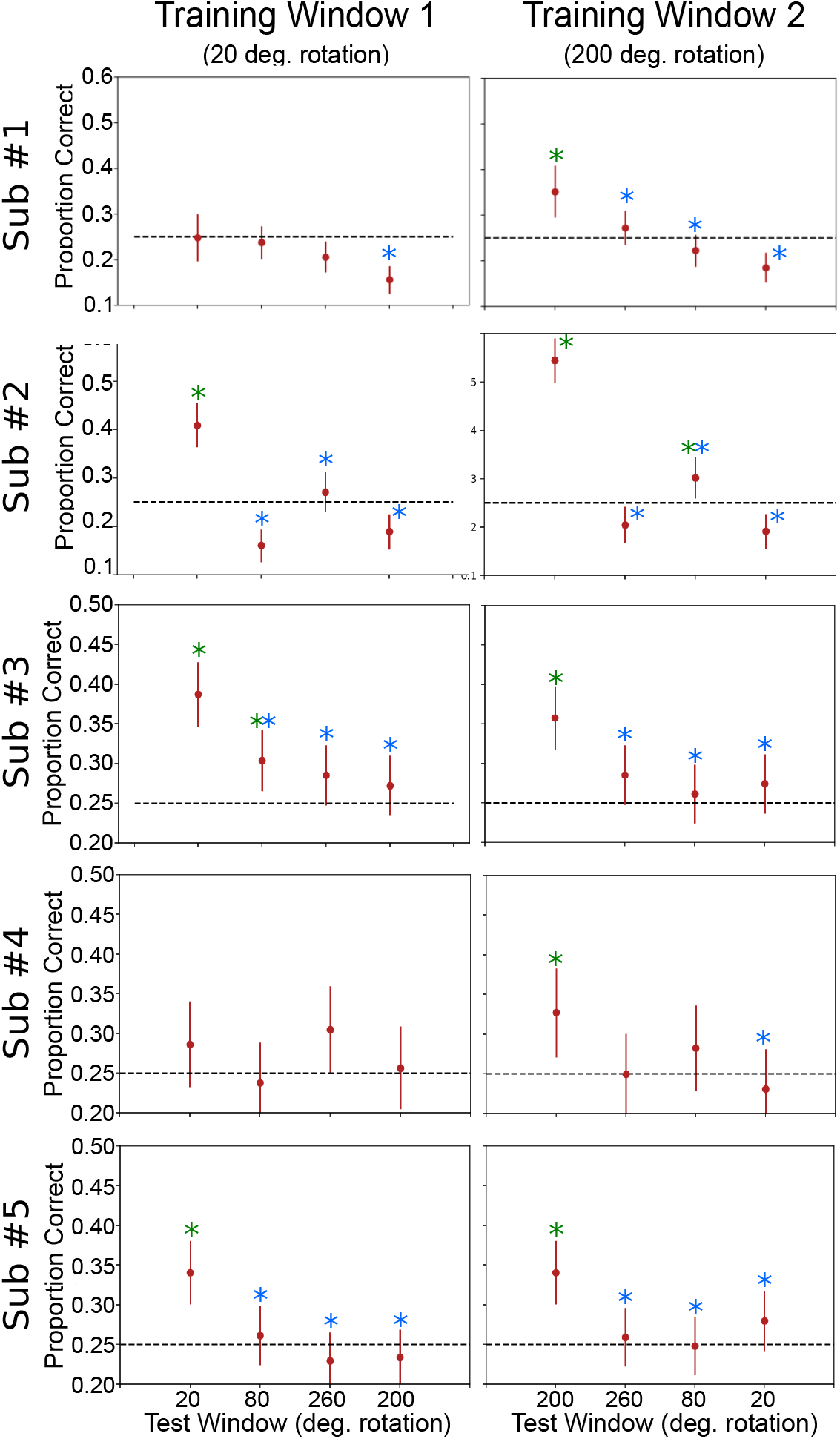
Classification accuracy results with test data for the decoding of orientation. Each row represents a complete analysis for a single subject. Columns represent different levels of the context dimension (spatial position) held fixed during training. For example, the leftmost cell for subject 1 displays accuracy results from a decoder that was trained to discriminate grating orientations, while holding the spatial position of the window constant at 20° of rotation. The same decoder was then tested with independent testing data in which the window was held at all spatial positions (20°, 80°, 260°, and 200° of rotation). A green asterisk represents accuracy values significantly above chance (dotted line), leading to the conclusion of invariance from the cross-classification test. As predicted, the cross classification test is susceptible to generating false positives, concluding that orientation is encoded invariantly from spatial position in the V1 of subjects 2 and 3. A blue asterisk represents that accuracy values significantly drop from the value observed at the trained window, leading to the conclusion of no invariance from the classification accuracy invariance test. The test is successful in finding evidence against invariance in the data of each participant.

### Jointly Testing Against Specificity and Invariance Leads To Valid Conclusions

The results from the previous section showed that the cross-classification test, which tests against the null hypothesis of context-specificity, can lead to erroneous conclusions about invariance of representations. We next aimed to show that the addition of tests of invariance developed from neurocomputational theory [4] could solve such issues and lead to more valid conclusions about the underlying code. Here, we apply two of these tests on our data set: the classification accuracy invariance test and the decoding separability test. In contrast to the cross-classification test, both of these theoretically-driven tests try to detect failures of invariance as opposed to providing evidence for invariance.

The classification accuracy invariance test defines invariance as the case where the probability of correct classification is exactly the same across all contexts. With invariance being the null hypothesis, the test is sensitive to any drop in the classifier’s performance across different levels of the context dimension. We implemented the classification accuracy invariance test by applying an omnibus Chi-Square test on the accuracy estimates from the linear decoder (i.e., testing whether all proportions are the same or some of them are different). Then, we performed pairwise comparisons between accuracy at the training level and each non-training level of the context dimension.

The decoding separability test, unlike the previous two tests, does not make use of classification accuracy estimates. Instead, it directly relies on certain properties of the decoding probability distributions for individual stimuli. That is, linear classifiers like the one used here perform classification of a new data point by computing a decision variable *z*, representing the distance of the data point from the classifier’s hyperplane separating two classes. When the decision variable is larger than some criterion value (usually zero), the output is one class, whereas when the decision variable is smaller than the criterion the output is the other class. Instead of comparing simple accuracy estimates, the decoding separability test compares the full distributions of such decision variables, or *decoding distributions*.

This test followed the same steps and rationale as the classification accuracy invariance test presented earlier, but instead of computing accuracies and testing their differences, we obtained decision variables from the trained classifier, and used those to estimate decoding distributions using kernel density estimation. For each pair of stimuli differing in the context dimension (e.g., 0° and 45° grating orientation, when the decoded variable was spatial position) we computed the distance between decoding distributions using a discretized *L*1 metric, which corresponds to the area highlighted in yellow in Figure 15. Then, we summed a number of such *L*1 metrics across values of the decoded dimension (e.g., the two spatial windows, when the decoded variable was spatial position), which produced an 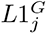 statistic (see Equation 3). Simply put, while a single *L*1 metric is analogous to the accuracy of the classifier for a single decoded label, the 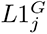 statistic is analogous to the overall decoding accuracy across all labels. The only difference is that 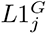 measures distances between decoding distributions, rather than accuracies. We performed a permutation test to determine whether the observed 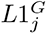 statistic was higher than expected by chance under the null hypothesis of invariance; a positive result on this test gives evidence against neural invariance for the given comparison. Also, we must note that, in theory, the decoding separability test should provide more information about (and be more sensitive to) such violations than the decoding accuracy invariance test (see 4).

As before, we first applied the invariance tests to decoding results from the spatial position classification. Results from the classification accuracy invariance are shown in Figure 5, and results from the decoding separability test are shown in Figure 7. The specific values obtained from the two tests are reported in Tables 2 and 3 of the Supplementary Material. The classification accuracy invariance test (Figure 5) did not find evidence against invariance in any of the subjects. However, in line with theoretical predictions, the decoding separability test (Figure 5) was much more sensitive to evidence against invariance present in the data. The test found failures of invariance in many cases where accuracy-based tests either found false positives (i.e., cross-classification) or failed to detect failures of invariance (i.e., classification accuracy invariance; see Figure 5). Overall, we found that the decoding separability test detected failures of invariance in the data of all five participants (17 out of 20 analyses).

**Figure 7:**
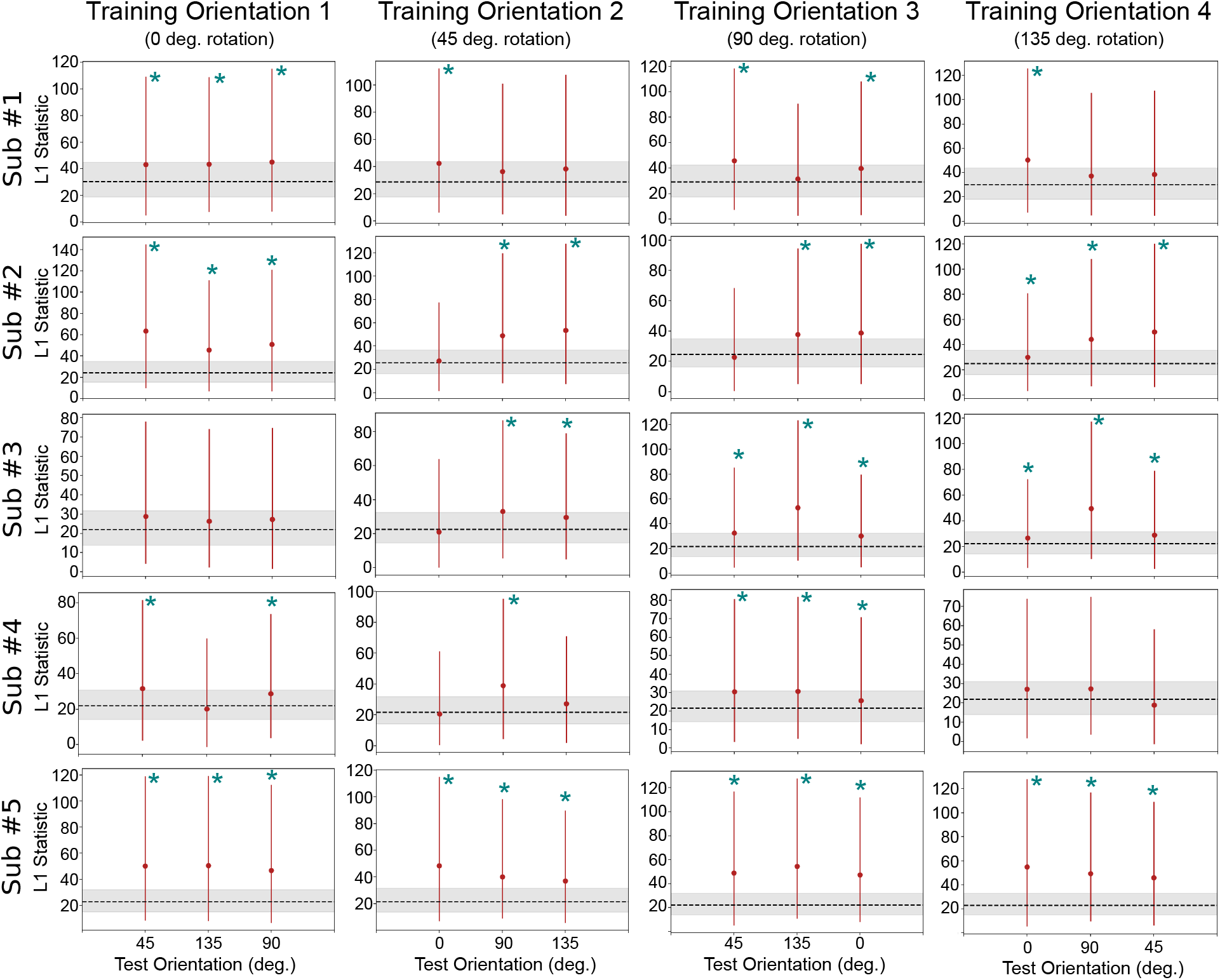
Decoding separability test results with spatial position as the target dimension. Each row represents a complete analysis for a single subject. Columns represent different levels of the context dimension (orientation) during training. The *y*-axis shows the 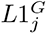 statistic, which quantifies the magnitude of violations of decoding separability. Bars represent 90% bootstrap confidence intervals on the 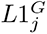 statistic. The dotted line and surrounded gray area represent the expected value and 90% bootstrap confidence interval for the 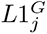 statistic when no differences exist between two distributions. Teal asterisks represent significant failures of invariance according to a permutation test. The test successfully found evidence for the absence of invariance in 18 out of 20 analyses.

From these results, it is apparent that the decoding separability test is sensitive to failures of invariance known to exist in the underlying neural code, even when decoding accuracy seems to suggest perfect invariance (see Figure 5). These results serve as an empirical validation of the decoding separability test, which was developed directly from theory [4]. In addition, these results show the value of testing against invariance, in addition to testing against specificity, to reach valid conclusions about the invariance or specificity of underlying neural representations. Performing both tests and following the guidelines in Table 1, results are inconclusive about whether encoding of spatial position in V1 is invariant or specific to orientation. This conservative conclusion is far better than the invalid conclusion that one would reach by performing the cross-classification test by itself; namely, that encoding of spatial position in V1 is invariant to orientation.

Next, we applied the invariance test to decoding results from the orientation classification. Results from the classification accuracy invariance test are shown in Figure 6, and results from the decoding separability test are shown in Figure 8. The specific values obtained from the two tests are reported in Tables 2 and 3 of the Supplementary Material. The classification accuracy invariance test (Figure 6) was much more sensitive to failures of invariance in this analysis. Failures of invariance were detected in every case where the classifier successfully decoded orientations above chance levels in the training window. Interestingly, failures of invariance were also detected in cases where the classifier did not successfully decode orientation above chance. This is counterintuitive, but expected from a theoretical point of view [see 4], which suggests that a decoder does not have to perform accurately or be optimal in any way to be able to detect failures of invariance. Contrary to our expectations, in this analysis the decoding separability test detected failures of invariance less frequently than the classification accuracy invariance test (see Figure 8). The decoding separability test detected failures of invariance in the data of four out of five participants (eight out of ten analyses).

**Figure 8:**
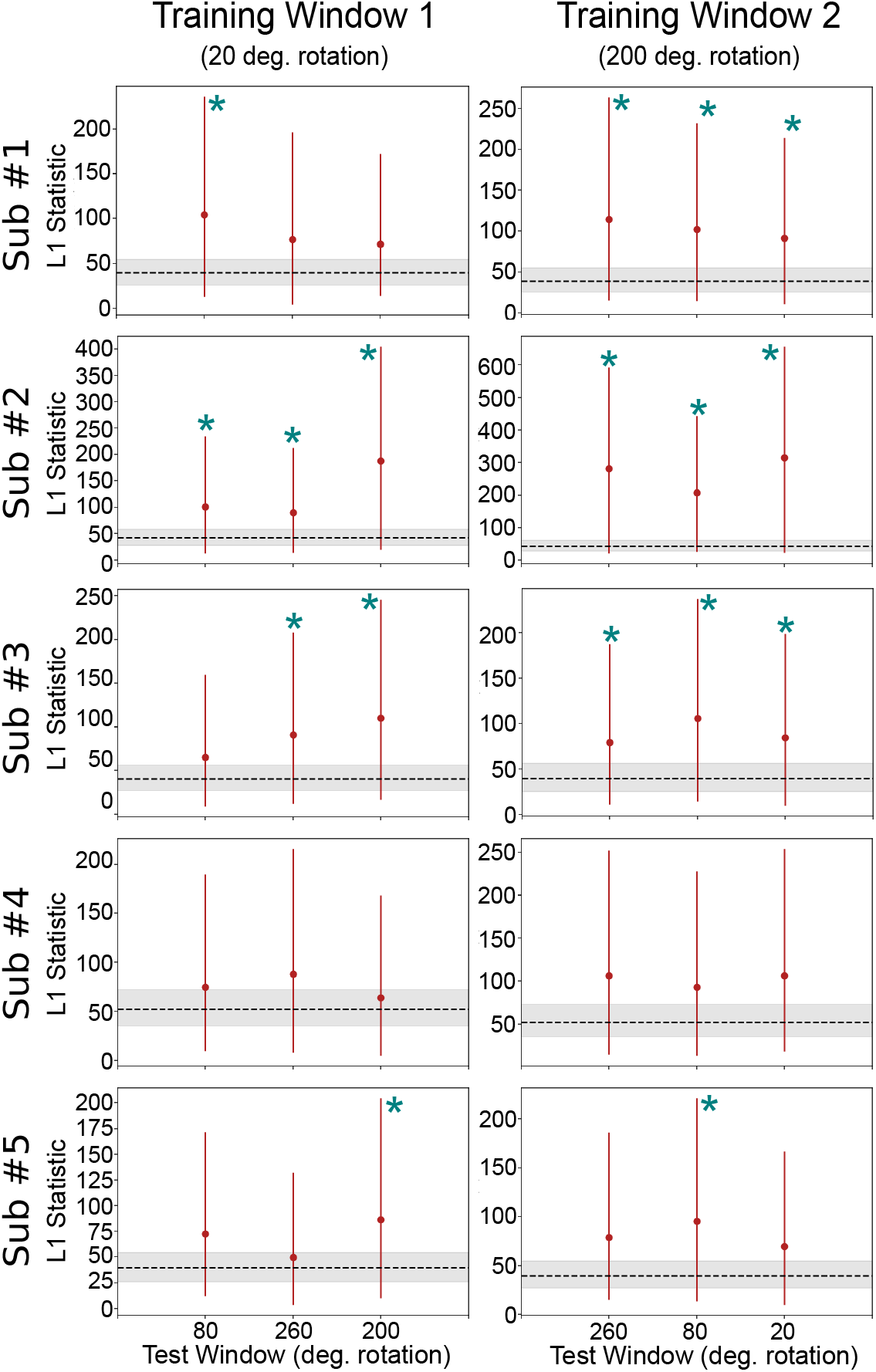
Decoding separability test results with orientation as the target dimension. Each row represents a complete analysis for a single subject. Columns represent different levels of the context dimension (spatial position) during training. The *y*-axis shows the 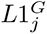 statistic, which quantifies the magnitude of violations of decoding separability. Bars represent 90% bootstrap confidence intervals on the 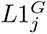 statistic. The dotted line and surrounded gray area represent the expected value and 90% bootstrap confidence interval for the 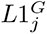 statistic when no differences exist between two distributions. Teal asterisks represent significant failures of invariance according to a permutation test. The test successfully found evidence for the absence of invariance in 8 out of 10 analyses.

In comparison to the classification accuracy invariance test, the decoding separability test appears to be more sensitive to detecting failures of invariance in cases where the decoder’s performance reaches ceiling levels (Figure 7). However, when classification accuracy is well below ceiling levels, as in decoding of orientation, the test seems less sensitive than classification accuracy invariance, perhaps due to a lower statistical power of the permutation test involved.

As was the case with decoding of spatial position, in this second analysis we also see the value of testing against invariance. While the results of the cross-classification test suggested invariant representations in subjects #2 and #3 (see Figure 6), such results are inconclusive when interpreted in the context of tests of invariance, which suggest context-specific representations.

### Simulation and Theoretical Results

The empirical results described in the preceding section clearly support the hypotheses that tests aimed at providing evidence for invariance, such as cross-classification, are prone to false positives, and that jointly performing tests against the nulls of invariance and specificity allows one to reach more precise and valid conclusions about the underlying representations.

However, there are issues with experimental work that motivated us to further evaluate our hypotheses through simulation work. In particular, experimental work does not allow full control of the underlying neural representations. In our study, we assumed that encoding of spatial position was specific to orientation, and vice-versa, but it is unlikely that the true encoding of these variables in V1 is completely context-specific. For example, there is evidence that a minority of neurons in V1 are invariant to orientation [21]. This means that encoding of spatial position is best characterized as context-sensitive, but a critical reader could interpret this as evidence for tolerance. Simulation work provides complete control over the representations under study, which can be made to be fully context-specific, without any degree of tolerance to changes in context. The relevant question is: Does cross-classification lead to conclusions of false-positive invariance under such circumstances? If yes: Can tests against invariance lead to more valid conclusions?

Another issue with experimental results is that they can be difficult to generalize. A critical reader could argue that issues with tests of context-specificity like cross-classification are restricted to special cases, and not general as suggested by theory. Again, simulation and theoretical work allows one to provide results that are more general.

### Simulation 1: False Positive Invariance Resulting From Features Of The Measurement Model

The empirical results presented in the previous section clearly show that the cross-classification test can generate false positives. Representations of orientation and spatial position in V1 can be categorized as context-sensitive at most (see Figure 1), but cross-classification can lead to conclusions of tolerance or invariance. Yet, some researchers might argue that they use the cross-classification test to detect *any* level of context-tolerance; that is, whether representations fall anywhere to the left of context-specificity in the continuum shown in Figure 1. In other words, some researchers might classify context-sensitivity as a form of tolerance, or partial invariance.

In theory, even a completely context-specific code could produce false conclusions of invariance in neuroimaging decoding studies, due to the transformation and mixing of neural responses from different populations that occurs at each voxel (see Figure 3). To provide evidence for such a general claim, we resort to simulation and theoretical work (for details on the models and procedures used in the simulations, see *Simulations* in the *Materials and Methods* section). We study a case of complete context-specificity in which it cannot be claimed that any amount of tolerance exists in the neural representations.

To create such a model, we started by defining two sets of encoding models, corresponding to two levels of the context dimension. In context 1, the target dimension was encoded through neural channels with homogeneous features (i.e., evenly spaced position, same maximum activity, same width), as shown at the top of Figure 18. In context 2, the target dimension was encoded through neural channels with completely randomized features, which is exemplified at the bottom of Figure 18. Then, we produced false positive invariance by optimizing the weights of the measurement model such that the voxel-wise activity values were similar across the two levels of the context dimension (Figure 18). Finally, we sampled data from both models and used them as input to a linear SVM classifier. As in the preceding empirical analyses, the decoder was trained on data from the first level model and tested on independent data from both the first and second level models (Figure 19). This entire procedure was repeated 200 times per simulation run, and we present the average results across simulations. We performed twenty simulation runs, where we gradually increased the measurement noise in each voxel (standard deviation going from 1 to 20, in steps of 1).

Figure 9a shows the decoding accuracy results from this simulation. The most important values are represented by the blue curves, which represent performance of the classifier in the non-trained level of the context dimension. Whenever accuracy is above chance, represented by the dotted line, the cross-classification test leads to a conclusion of invariance in a situation where no invariance exists (i.e., false positive invariance). The cross-classification accuracy score was much higher than chance across all levels of noise, even as measurement noise was drastically increased. The red line in Figure 9b shows the proportion of false positives for the cross-classification test, which consistently remained above the nominal *α* = .05, represented by the dotted line, across all levels of noise that produced above-chance decoding. Only when decoding accuracy drops to chance levels (a case where the test would not be applied in an empirical setting) the cross-classification test stops producing false positives.

**Figure 9:**
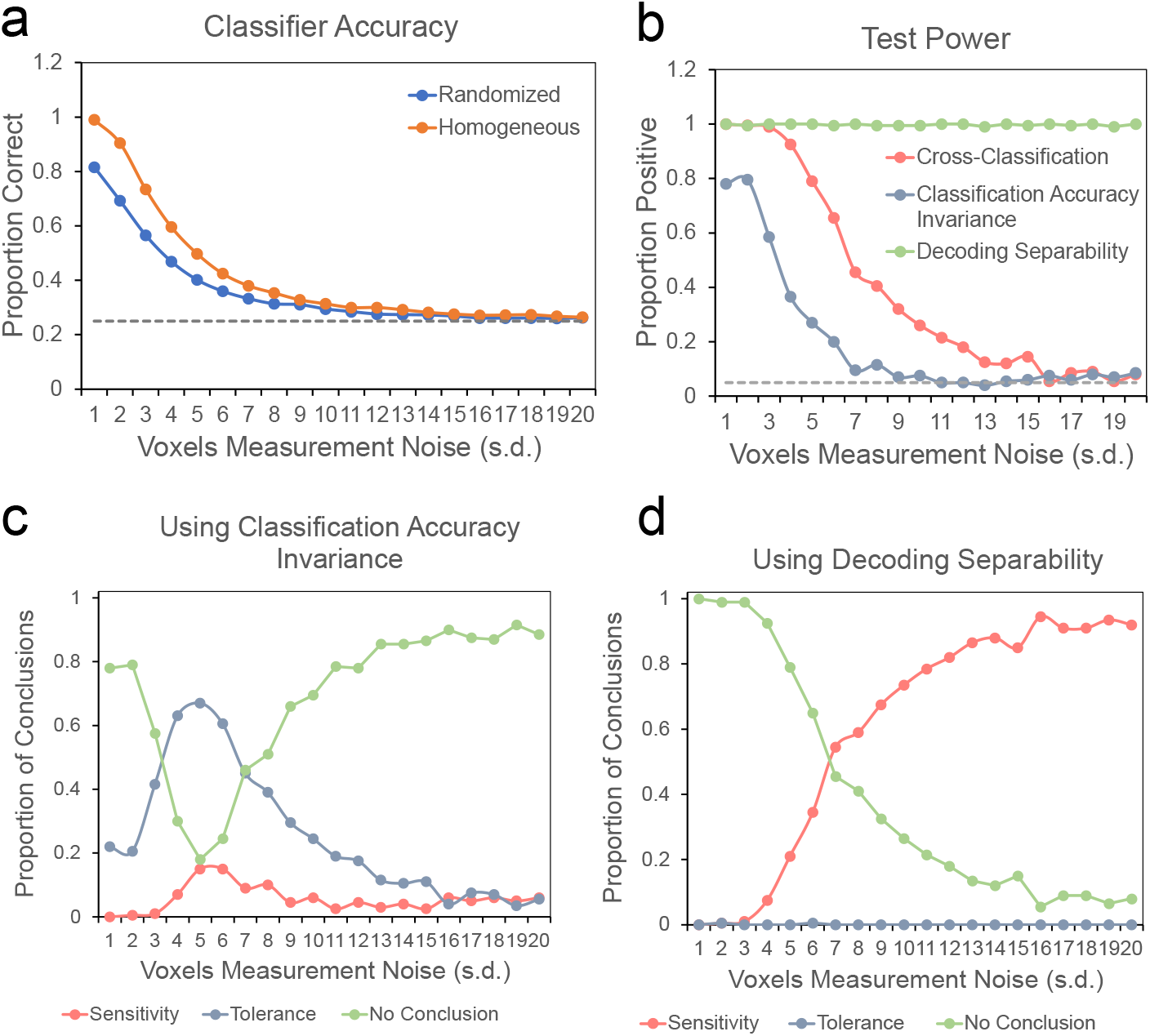
Decoding results from simulation 1. (a) Classifier accuracy scores for model-generated data from both levels of the context dimension. The *y*-axis represents accuracy scores, while the *x*-axis represents level of measurement noise (in units of standard deviation). (b) Proportion of positive tests of each type. The *y* - axis represents proportion of positives, and *x*-axis represents measurement noise. At all levels of measurement noise producing classifier accuracies above chance, the proportion of false positive cross-classification tests remains higher that the accepted 5% threshold (dotted line). (c-d) Proportion of each type of conclusion in Table 1 (specificity/sensitivity in red, invariance/tolerance in blue, and no conclusion in green) reached from jointly testing against specificity and invariance. The left panel (c) shows conclusions reached by using the classification accuracy invariance test against invariance, and the right panel (d) shows conclusions reached by using the decoding separability test against invariance. In both cases, the cross-classification test is used against specificity.

These results suggest that a suitable selection of measurement model is sufficient for inducing false positives in the cross-classification test, even when the underlying encoding distributions themselves show absolutely no tolerance. The next question is whether using additional tests against the null of invariance can lead to more valid conclusions.

The blue line in 9b shows the proportion of tests correctly rejecting the null of invariance for the classification accuracy invariance test. The test is very sensitive to measurement noise, having good power (about 80%) only at the smallest levels of measurement noise. Figure 9c shows the proportion of each type of conclusion in Table 1 (specificity/sensitivity in red, invariance/tolerance in blue, and no conclusion in green) reached from jointly testing against specificity and invariance, by using the cross-classification and classification accuracy invariance tests, respectively. This strategy does lead to more valid conclusions at either low or high levels of noise, but at intermediate levels the strategy fails and produces a high proportion of conclusions for tolerance. Note that these intermediate levels of noise produce decoding accuracy around 40%-60%, which are realistic values for a four-alternative classification task.

The green line in Figure 9b shows the proportion of tests correctly rejecting the null of invariance for the decoding separability test. The first notable result is the high sensitivity of the decoding separability test to violations of invariance. At all levels of noise, the test detected such violations in almost all simulation runs. Note that the test is sensitive even when decoding accuracy has dropped to chance. All these features of the test are expected from the theory used to develop it [4]. Higher sensitivity than accuracy-based tests is expected because the test uses information from the full distribution of decision variables from the decoder. Robustness in the face of measurement noise is expected because although noise reduces high-frequency differences between distributions, it preserves differences at lower frequencies (see [4]). We must note that this simulation probably over-estimates the test’s sensitivity, as our experimental results showed that the test often misses significance in real data.

Figure 9d shows the proportion of each type of conclusion in Table 1 (specificity/sensitivity in red, invariance/tolerance in blue, and no conclusion in green) reached from jointly testing against specificity and invariance, by using the cross-classification and decoding separability tests, respectively. In this case, invalid conclusions of tolerance are never reached. Counterintuitively, valid conclusions of sensitivity increase over inconclusive results as noise increases. The reason is that cross-classification is more sensitive to noise than decoding separability.

Overall, the results from this simulation provide further evidence favoring our hypotheses, showing that cross-classification can lead to false positive conclusions of tolerance when absolutely no tolerance exists in the underlying neural code, and that the addition of tests against invariance leads to more valid conclusions. The results suggest that decoding separability should be preferred over classification accuracy invariance to test against invariance, as was expected from theory [4].

### Evaluating the Pervasiveness of the False Positive Invariance Problem

A critical reader might argue that the conditions leading to false positive invariance in the first simulation, namely the explicit selection of the measurement weights that produce similar voxel-wise activity patterns across levels of the context dimension, are unlikely to occur in real fMRI experiments. The true measurement process is not trained to make activity values similar across different levels of irrelevant dimensions. How pervasive is the false positive invariance problem uncovered in the first simulation? Here we show that, against intuition, the problem is quite pervasive.

In the standard encoding model used in our simulations, the mean response of neural channel *c* to stimulus *s_i_*, presented in context *j*, is given by a tuning function *f_jc_*(*s_i_*) (see subsection in Materials and Methods). We can collect the mean response of *N_c_* channels in a population response vector **f***_j_* (*s_i_*) = [*f_j_*_1_(*s_i_*)*, f_j_*_2_(*s_i_*)*, …f_jNc_* (*s_i_*)]. A number of stimulus values for the target dimension are presented in any experiment, indexed by *i* = 1, 2, …, *N_s_*. Without loss of generality, we can focus on an experiment with two stimulus contexts indexed by *j* = 1, 2, as in our simulation. The measured activity in voxel *k* to stimulus *s_i_* in the first context is equal to **f**_1_(*s_i_*)^T^**w***_k_*_1_, and in the second context is equal to **f**_2_(*s_i_*)^T^**w***_k_*_2_. The measurement vectors **w***_k_*_1_ and **w***_k_*_2_ produce invariance in voxel *k* when they produce the same mean activity value:

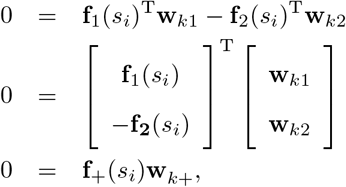

where **f**_+_(*s_i_*) is a row vector of concatenated mean population responses, and **w**_+_ is a column vector of concatenated weights.

If we collect the vectors **f**_+_(*s_i_*) in response to the experimental stimuli in a matrix:

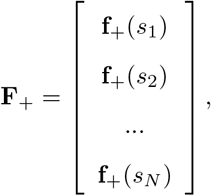

we get a set of homogeneous equations that can be solved for **w***_k_*_+_ *>* 0:

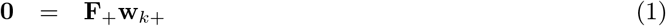

A measurement model produces false positive invariance when **w***_k_*_+_ ≠ **0** is a solution of this equation for all voxels *k*. Another way to see this equation is that **w***_k_*_+_ corresponds to the nullspace of matrix **F**_+_. The *nullity-rank theorem* tells us that the dimensionality of this nullspace, or nullity, equals the number of columns in **F**_+_ (i.e., the total number of channels in the model) minus its rank. The nullity gives us information about the size of the subspace of measurement models **w***_k_*_+_ that produce false positive invariance. When the only solution for Equation 1 is the trivial solution **w***_kc_* = **0**, the nullity of **F**_+_ is zero. In this case, constraints in the encoding model and experimental design, summarized in **F**_+_, are such that there is no measurement model that can produce false positive invariance. This is the *only* case in which we would not have to worry about false positive invariance, but it has been the default assumption of researchers applying the cross-classification test in the literature. Note also that this analysis is only concerned with *strict invariance* and not with *tolerance*; even when false positive invariance cannot be produced by a measurement model, false positive tolerance may still be possible.

We are now in a good position to evaluate the pervasiveness of false positive invariance in the encoding scenario posed by our first simulation. We created encoding models just as indicated for simulation 1 (see Figure 18), each time with a different number of stimuli and neural channels. The number of neural channels was varied from 5 to 30 in steps of 5, and the number of stimuli was varied from 2 to 20 in steps of 2. For each combination of neural channels and stimuli, we created 200 different encoding models, and computed the nullity of the mean population response matrix **F**_+_. As indicated earlier, the nullity represents the dimensionality of the subspace of measurement models that would produce false positive invariance. To ease comparison, Figure 10 shows the nullity divided by the dimensionality of the measurement model, or proportion nullity. This represents the proportion of the measurement space (in terms of dimensionality) that would produce false positive invariance. We found that there was no variability of results across the 200 sampled models, so Figure 10 shows the unique value of proportion nullity found in each case.

**Figure 10:**
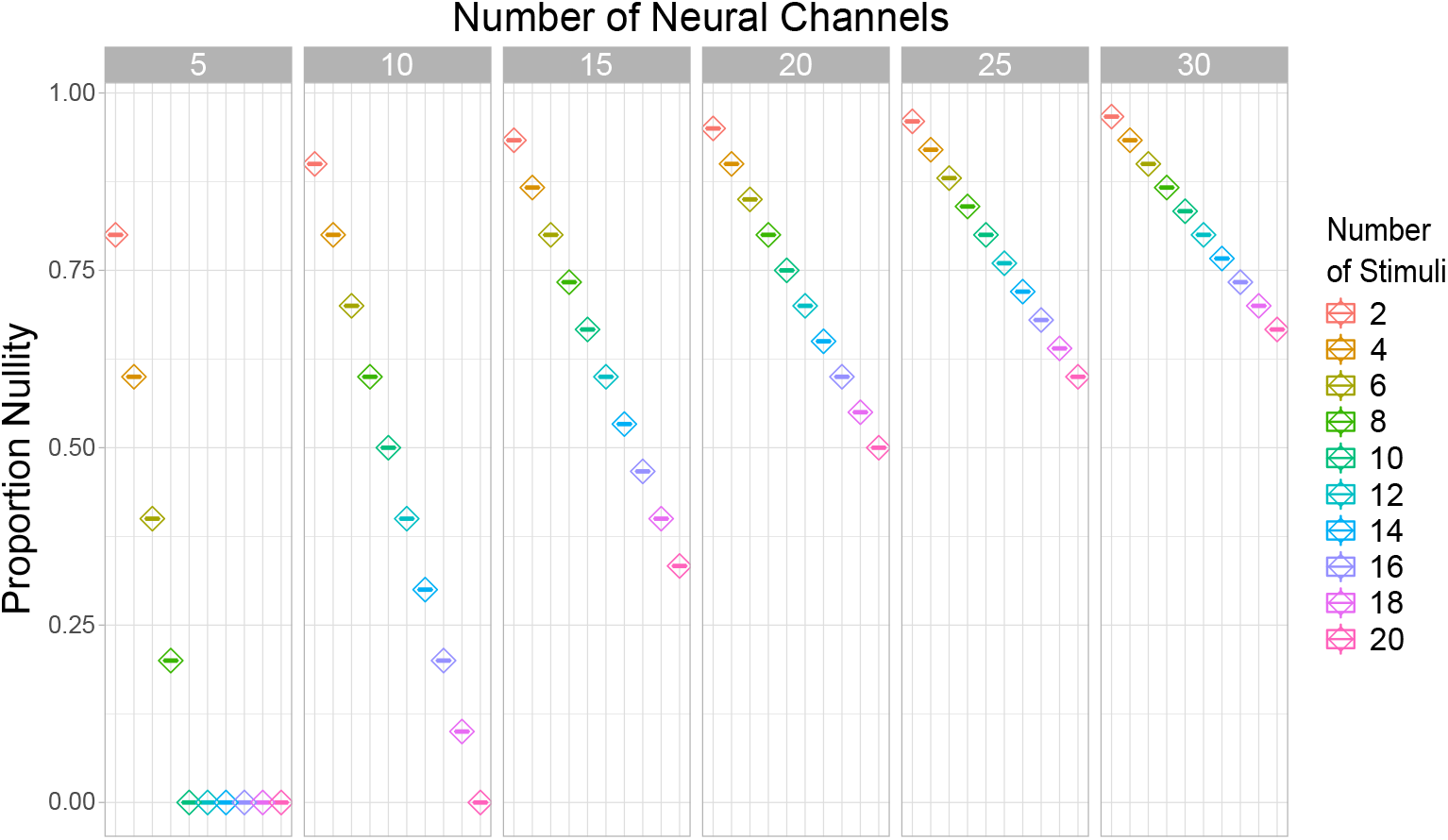
Pervasiveness of the problem of false positive invariance for the extreme case of context-specificity studied in simulation 1. Proportion nullity represents the proportion of all dimensions in the measurement space that would produce false positive invariance, and therefore the size of the false positive invariance problem. The values reported were always the same for a given combination of number of neural channels and number of stimuli, across 200 randomly sampled encoding models.

One can easily see from Figure 10 that the scenario posed in our first simulation is far from rare. On the contrary, with ten channels and four stimuli, as we used in that simulation, the proportion nullity is 0.8, meaning that the large majority of the possible measurement models will lead to false positive invariance. This result was not idiosyncratic to the parameters chosen for our simulation, with proportion nullity in general being quite high. The exception was a combination of a high number of stimuli and low number of channels, which is rare in experiments reported in the literature. Most neuroimaging studies using cross-classification to study invariance have presented 2-4 stimuli, a case in which the proportion nullity is at least 0.6, and in most cases above 0.8.

We must remind the reader that we are studying here an extreme case of context-specificity, under the assumption of no measurement noise, and an extreme case of false positive invariance, rather than tolerance. For these reasons, we can consider our results a lower bound on the size of the false positive invariance problem. More realistic scenarios involving context-sensitivity, high measurement noise, or evaluation of tolerance rather than strict invariance can all be expected to worsen the problem beyond what is shown in our results.

We see two clear trends in Figure 10. First, proportion nullity –and therefore, the problem of false positive invariance– drops linearly with number of stimuli included in the study. Experimenters can reduce the risk of false positive invariance by increasing the number of stimulus levels for the target dimension. Second, proportion nullity increases in a negatively accelerated fashion with increments in the number of neural channels. The number of neural channels represents our assumption of how many unique neural tuning functions underlie the data or, in other words, how well-covered is the stimulus dimension by the encoding neural population. In realistic scenarios, this value will be much higher than any of those shown in Figure 10. However, it is common to find applications of the standard encoding model in computational neuroimaging that assume 6-15 channels [e.g., 22, 23, 24, 25].

### Simulation 2: False Positive Invariance Resulting From Similarly Tuned Neural Subpopulations Across Contexts

A critical reader may again argue against the results just presented, indicating that although the space of possible measurement models leading to false positive invariance is large in most published studies, most of those models would never be observed in nature. Only a small proportion of all possible measurement models might be truly at play in neuroimaging studies, and those could be contained within the space of models for which false positive invariance is not an issue. Although this is an extremely optimistic position, and we think that it would be unwise for scientists to take it, we would like to strengthen our conclusions by studying a realistic encoding scenario, likely to be implemented in the brain.

There are many known cases in which neurons that are sensitive to a particular stimulus feature are spatially clustered at sub-millimeter scales. In those cases, while there is spatially distributed information about stimulus features, this information is not immediately visible at the typical resolution of an fMRI study. For example, V1 neurons that are sensitive to the same spatial frequency, color, ocular dominance, and orientation all cluster at the sub-millimeter scale [26, 27, 28, 29]. Although advances in high-field fMRI can in some cases uncover such sub-millimeter organization [e.g., 30], information can also be spatially distributed without any clustering (e.g., “salt-and-pepper” codes; see [31, 32]), at scales that are unlikely to be reached with fMRI at even higher field strengths than those currently available [33, 34].

In cases such as these, across voxels we would expect to find relatively homogeneous distributions of selectivities. Our ability to use voxel-level decoding to detect whether and how features are encoded depends critically on small random variations in mixing; that is, in the proportion of each type of neuron present within each voxel. Indeed, small differences in mixing across voxels is a mechanism proposed to underlie decoding of orientation information from V1 [35, 36, 37, 38, 39], like that shown in our experimental study.

This sub-voxel distribution of information, which may underlie the success of many fMRI decoding studies, can also easily lead to false-positive invariance when the cross-classification test (or other tests of the null of specificity) is used in isolation. Small differences in mixing might be enough to promote above-chance decoding of a stimulus feature, because decoding algorithms are specifically trained to detect differences in the target feature. On the other hand, decoding algorithms are not trained to detect changes in context. Any small differences in mixing that might provide information about context-specificity would be lost, and the decoding algorithm would be very likely to generalize performance across changes in stimulus context. We find an example of this in our own experiment. There, classification of spatial position generalized perfectly across changes in grating orientation, as shown in Figure 5, despite the fact that the voxels contained information about differences in orientation, as determined by above-chance decoding of that dimension (see Figure 6).

In the present simulation, we wanted to study the sensitivity of different fMRI decoding tests to changes in mixing carrying information about context-sensitivity. With this goal in mind, we created a model in which a target dimension is encoded in a completely context-specific manner, with one subpopulation of neurons responding whenever the context dimension is at level 1, and a different subpopulation of neurons responding whenever the context dimension is at level 2. Both subpopulations were modeled using a standard homogeneous encoding model (see above), but note that this similarity in tuning functions is not equivalent to invariance, as each channel responded *only* at one of the levels of the context dimension. In other words, our simulation assumes that populations encoding the target dimension are completely separated across levels of the context dimension, but they encode the target dimension in a similar way (just as neurons in Figure 4 have two selectivity types across levels of the context dimension). As before, we report the averaged results from 200 simulations in each run. Measurement noise was set to a fixed level across simulations (s.d.=5, which in our previous simulation produced accuracies around 40%-50%, see Figure 9). In each simulation run, we increased the difference in the measurement models for the two levels of the context dimension, by adding random noise to weights of the measurement model as illustrated in Figure 20. The standard deviation of the weight noise was gradually increased from 0.05 to 0.5 (i.e., from 0.5 to 5 times the average weight value), in steps of 0.05. That is, in the final models the contribution of each neuron type (e.g., neurons selective to a value of 0 in the target dimension) was widely different across levels of the context dimension.

The results from this simulation are shown in Figure 11. Panel a shows the accuracy of the classifier tested in the original training context (red line) and in the changed context (i.e., cross-classification performance; blue line). It can be seen that the cross-classification test is sensitive to mixing variations, as accuracy drops with increments in weight changes with context. However, accuracy remains well above chance even for the largest weight changes. Figure 20b shows the proportion of positive tests as a function of the magnitude of random weight changes (in standard deviations). The cross-classification test consistently showed false positives at a rate much higher than the nominal 5%. High levels of false-positive invariance were present even when the weight noise standard deviation was five times as large as the average weight values. These results suggest that, when two *completely separate* neural populations use similar codes to represent a target dimension across levels of an context dimension, false positive invariance is likely to be found not only with the small variations in mixing that one would usually expect from fMRI studies, but from very large variations in mixing.

**Figure 11:**
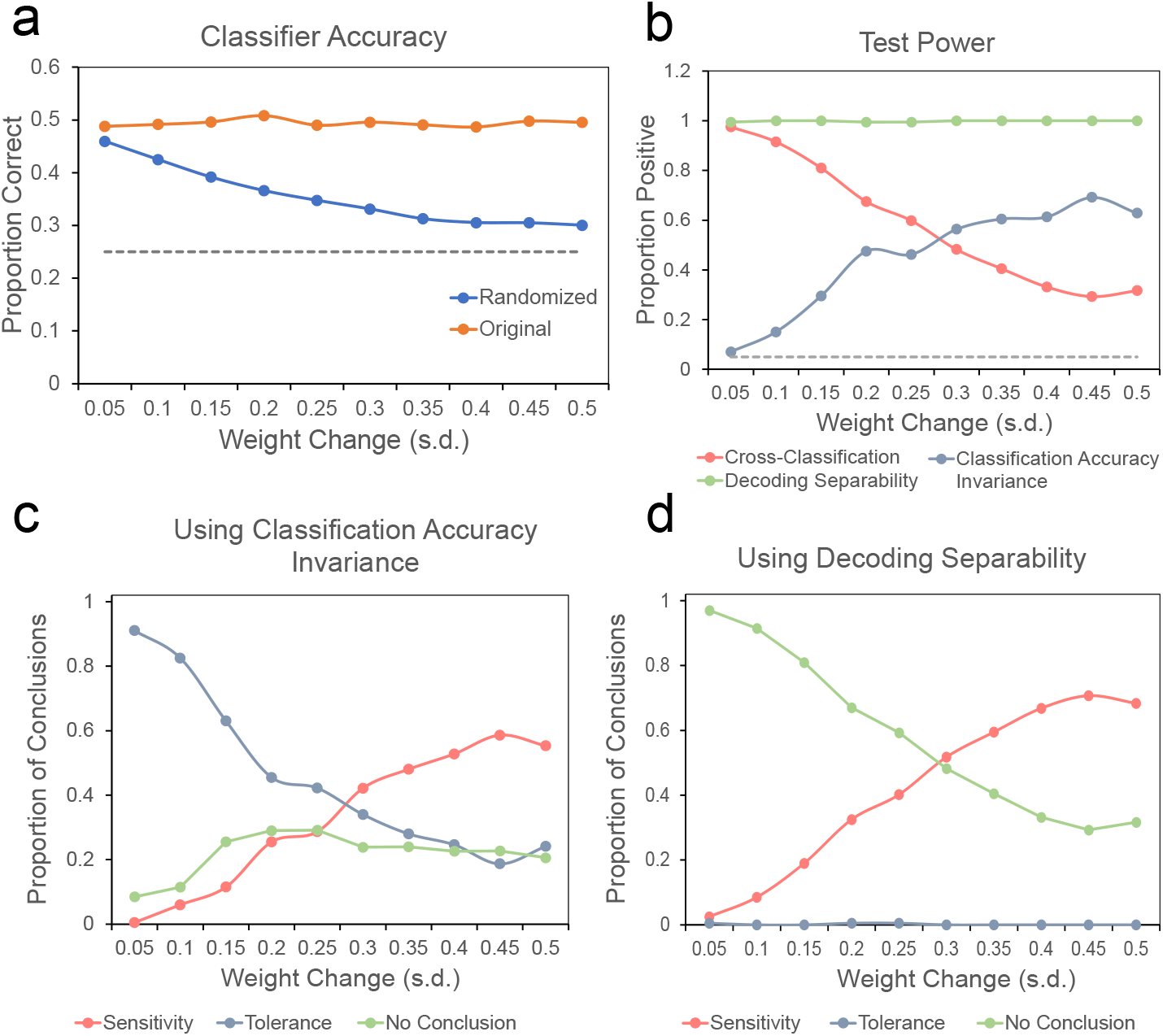
Decoding results from simulation 2. (a) Classifier accuracy scores for model-generated data from both levels of the context dimension. The *y*-axis represents accuracy scores, while the *x*-axis represents magnitude of noise added to measurement weights for the second level model. Note that classifier accuracy never dips below chance levels (dotted line) for either model. (b) Proportion of positive tests of each type. The *y*-axis represents proportion of positives, and *x*-axis represents measurement noise. At all levels of measurement noise producing classifier accuracies above chance, the proportion of false positive cross-classification tests remains well above the accepted 5% threshold (dotted line) at noise levels with a standard deviation five times the average value of the original weights. (c-d) Proportion of each type of conclusion in Table 1 (specificity/sensitivity in red, invariance/tolerance in blue, and no conclusion in green) reached from jointly testing against specificity and invariance. The left panel (c) shows conclusions reached by using the classification accuracy invariance test against invariance, and the right panel (d) shows conclusions reached by using the decoding separability test against invariance. In both cases, the cross-classification test is used against specificity.

As before, the question now is whether this issue of false-positive invariance can be ameliorated by adding tests against the null of invariance. Using classification accuracy invariance, the results are not very promising. The blue line in Figure 20b shows the power of this test to reject the null of invariance, which starts near zero with very small variations in weights (or mixing) and is quite low (∼60% power) even at the largest weight variations. Figure 20c shows the proportion of each type of conclusion in Table 1 (specificity/sensitivity in red, invariance/tolerance in blue, and no conclusion in green) reached from jointly testing against specificity and invariance, by using the cross-classification and classification accuracy invariance tests, respectively. First, the addition of classification accuracy invariance does improve the validity of conclusions. Comparing the red curve in Figure 20b against the blue curve in Figure 20c shows that the latter drops more steeply with size of weight changes. On the other hand, using cross-classification and classification accuracy invariance together still leads to a false positive rate above 5% across all the values of weight change simulated.

On the other hand, using decoding separability the results are much better. The green line in Figure 20b shows that the power of this test to reject the null of invariance is near 100% across all levels of weight change. That is, the test is sensitive to even small changes in mixing resulting from changes in context. As explained before, this high power is a consequence of the test using the whole distribution of decision variables from the decoder, rather than only binary classification decisions. Figure 20d shows the proportion of each type of conclusion in Table 1 (specificity/sensitivity in red, invariance/tolerance in blue, and no conclusion in green) reached from jointly testing against specificity and invariance, by using the cross-classification and decoding separability tests, respectively. First, the test has a higher power than classification accuracy invariance to reach the correct conclusion of context-sensitivity. More importantly, at low values of mixing, the test leads to inconclusive results rather than to the incorrect conclusion of invariance.

Overall, this simulation confirms our previous conclusion and theoretical expectation that supplementing the cross-classification test with a test against the null of invariance increases the validity of conclusions about the underlying codes, and that decoding separability is superior to classification accuracy invariance for that goal. We have shown that this is the case in the realistic scenario in which two different populations encode the target dimension in a similar manner, and the fact that both populations are separated can be inferred only from small differences in their relative contribution to voxel activities. With such small differences in mixing (i.e., the smallest values of weight change in Figure 20), using cross-classification alone leads to a conclusion of false positive invariance almost 100% of the time, the addition of the classification accuracy invariance test slightly reduces the issue, and the addition of the decoding separability test eliminates it, at least in our simulation. The results were similar at very large differences in mixing (i.e., the largest values of weight change in Figure 20), where using cross-classification alone leads to a conclusion of false positive invariance about 30% of the time, the addition of the classification accuracy invariance test slightly reduces the issue, and the addition of the decoding separability test eliminates it, with the most likely conclusion being the ground truth of context-sensitive encoding. We must warn again, however, that the sensitivity of the decoding separability test is expected to be lower with experimental data, as it was in our own study. The main reason why decoding separability is so extremely powerful in our simulations is that they assumed the extreme case of completely context-specific codes.

## Discussion

Here, we have provided empirical and computational evidence supporting two insights about decoding tests of invariance reached with the help of neurocomputational theory [4]. First, that tests aimed at evaluating evidence against the null of context-specificity, and for the alternative of context-invariance, may be prone to false positives due to the way in which the underlying neural representations are transformed into measurements. Second, that jointly performing tests against the nulls of invariance and specificity allows one to reach more precise and valid conclusions about the underlying representations.

In the empirical study, we performed decoding of orientation and spatial position from fMRI activity patterns recorded in V1, a case in which properties of the underlying neural code are known. The cross-classification test gave strong evidence for the incorrect conclusion that, in V1, encoding of spatial position is tolerant/invariant to changes in orientation, as well as some evidence for the incorrect conclusion that orientation is tolerant/invariant to changes in spatial position. We found that the addition of theoretically-derived tests of invariance leads to more valid conclusions regarding the underlying code.

The results of two simulations strengthened the conclusions from the empirical study, by showing that they hold even in the extreme case of completely context-specific encoding. In the first simulation, we showed that cross-classification can lead to false positive conclusions of tolerance when absolutely no tolerance exists in the underlying neural code, and that the addition of tests against invariance leads to more valid conclusions. We also showed, through theoretical analysis and further simulations, that this problem is likely to be pervasive, rather than resulting from a hand-picked proof of concept. In our second simulation, we showed that the same results are found in simulations of realistic encoding scenarios.

Based on our empirical and computational results, we conclude that the cross-classification test can lead to invalid conclusions about the invariance of neural representations. Applying the test by itself should be avoided, and previous research using the test should be re-evaluated in light of our results. Instead, we propose to routinely test against the null of invariance whenever the cross-classification test is applied. Even if a researcher is unconvinced by the pervasiveness of the problem highlighted in our study and simulations, the cost of running these additional tests is extremely low.

As expected from theory, we found that the decoding separability test is sensitive to violations of invariance that cannot be captured by the classification accuracy invariance test. In particular, when decoding accuracy is near ceiling or floor values, only the decoding separability test can detect violations of invariance by relying on the more fine-grained information available in the full decoding probability distributions, rather than on the coarse information available in accuracy estimates. The reason behind this superiority is simple: the decoding separability test uses information from the full distribution of decoder decision variables, and much of this information is lost once that distribution is binarized for classification. A similar conclusion was reached by Walther et al. [40], who found that the reliability of continuous neural dissimilarity measures was higher than that of classification accuracies, and concluded that this was due to the loss of information inherent to the latter. We believe that focusing on full decoding distributions can help us to move from using decoding to test *whether* information is encoded in a particular area, to using decoding to test *how* information is encoded. Additional examples of this approach have linked uncertainty in decoding distributions to behavior [41], and have correlated the variability in decoding distributions to behavioral responses [42].

A drawback of the decoding separability test is that its application requires a large number of trials per participant. Precise kernel density estimates (and more generally, precise estimation of differences between two distributions), which are essential for accurate results from the decoding separability test, require large longitudinal datasets like the one used in this study [43]. With a small number of trials per stimulus, the classification accuracy invariance test might be a better choice.

Relatedly, the study of invariance could benefit from the development of a test against context-specificity to replace the cross-classification test, which is relatively insensitive due to its reliance on decoding accuracy. This new test should aim to evaluate the null hypothesis that the neural representation of a target stimulus property is completely different in two different contexts, showing non-overlapping distributions of neural activity. The development and validation of such a measure is not trivial, and we must leave it to future research. We at least know that a sensitive measure would rely on something different from the decoding distribution of the target variable, and therefore it would follow a different logic than the decoding separability test developed in previous work [4]. Figure 12 shows why this is the case. The two main axes represent measurements in two different voxels, and each ellipse represents the distribution of voxel activity patterns for a target stimulus property presented in two different contexts. It can be seen that the two distributions are completely non-overlapping in the multivariate space of voxel patterns. However, when the two distributions are projected onto the decoded variable they show a non-zero overlap, represented by the yellow rectangular area. Note how changing the direction of the decoded variable does not necessarily result in no overlap. Also, simply measuring the overlap at the voxel level does not solve the issue, because the measurement model may also artificially introduce overlap in the distributions that does not exist in the underlying neural representations. This is simply a case of the more general issue with measurement models inducing invariance in context-specific representations, presented in Figure 3b.

**Figure 12:**
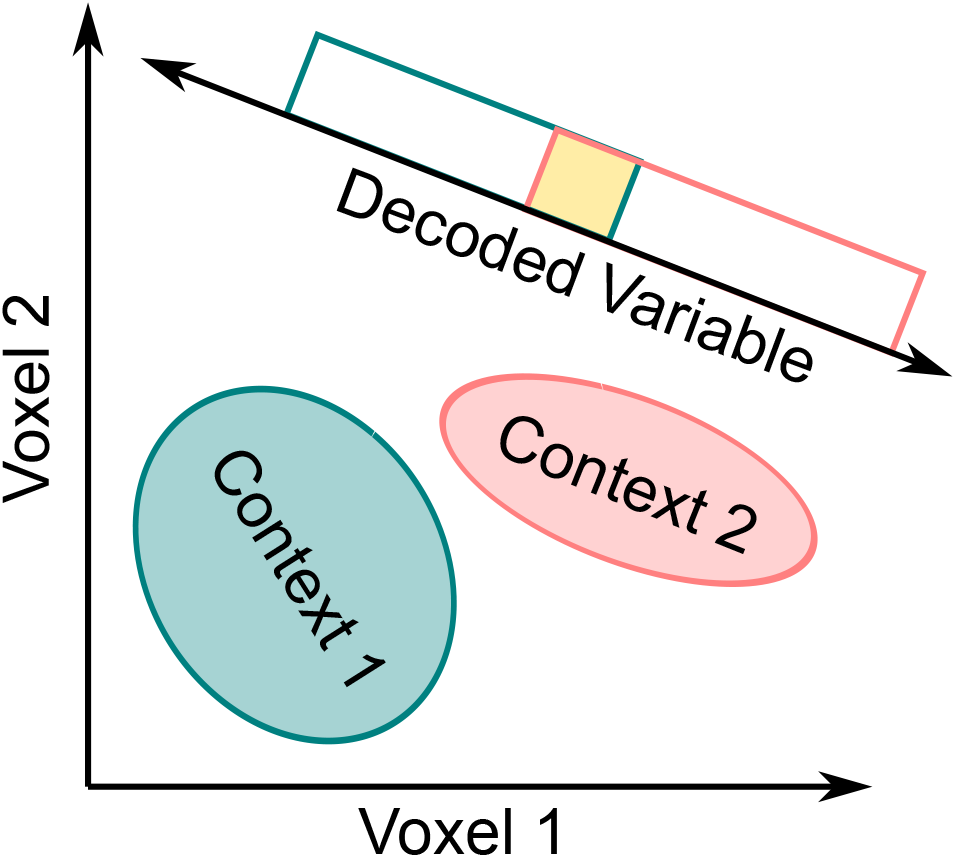
The decoding distributions along the decoded variable cannot be used to obtain a valid test of no overlap between neural representations across two contexts. The main axes represent measurements at the voxel level, and each ellipse represents the distribution of neural activity (after transformation by the measurement model) for a target stimulus property presented in two different contexts. The two distributions are completely non-overlapping at the level of the multivariate voxel patterns. However, when the two distributions are projected onto the decoded variable, they show a non-zero overlap represented by the yellow area.

Beyond the specific case of decoding tests of invariance, the present study shows the dangers of over-reliance on operational tests that have only face validity, particularly in the study of neural representation through indirect measures obtained through neuroimaging. Our study joins other recent reports in the literature [25, 44] in showing that the application of sophisticated data analysis tools can lead to the wrong conclusions when problems of identifiability (e.g., between neural and measurement factors) inherent to neuroimaging are not taken into account. We believe that theoretical and simulation work will play an important part in the future of neuroimaging, both to point out areas in which our methods might run into issues, as well as showing us potential solutions.

## Materials and Methods

### Participants

Five healthy volunteers (ages 19–27, three female) from Florida International University participated in the experiment; all had normal or corrected-to-normal vision. The study protocol was approved by Florida International University’s Institutional Review Board and by the Center for Imaging Science Steering Committee. All subjects gave written consent to experimental procedures before participating in the experiment.

### Stimuli

All stimuli were generated using Psychopy v.1.85.0 [45]. Images were displayed on a 40″ Nordic Neurolab LCD InroomViewing Device, placed at the rear entrance of the scanner bore. Subjects viewed the screen via an angled mirror attached to the head coil. Visual stimuli were full-contrast square-wave gratings with a spatial frequency of 1.5 cycles per degree of visual angle [similar to 18, 19, 46], a frequency known to drive V1 responses strongly [47], shown through a wedge-shaped aperture window that spanned from 1.5° to 10° of eccentricity and 100° of polar angle (Figure 13). The aperture window had four possible locations, starting at 20°, 80°, 200°, and 260° of rotation. The square-wave gratings were oriented in one of four angles for each trial: 0°, 45°, 90°, 135°. The phase of the gratings was randomly changed every 250ms, to reduce retinal adaptation and afterimages.

### Task and Procedures

To ensure that the data used to train a classifier in decoding analyses (see below) was independent from the data used to test the classifier and compute measures of performance, training trials and testing trials were presented on separate acquisition runs. Training and testing runs were identical in all aspects except one: the positions of the aperture window were restricted to 20° and 200° of rotation for the training runs, while testing runs included all four positions (Figure 13). During stimulus presentation, the phase of the grating was randomly shifted every 250 ms. The orientation of each grating was randomly chosen on each trial, while the spatial position of the window changed sequentially in a pre-determined manner. In training runs, the aperture window switched between 20° and 200° on every trial. In testing runs, the aperture window cycled through 20°, 200°, 80°, 260°, in that order. For both training and testing runs, each combination of spatial position (two or four levels) and orientation (four levels) was presented 35 times in a single acquisition session. Each subject went through 4 identical acquisition sessions to yield a total of 135 presentations of a given combination of orientation and spatial position (see all combinations in Figure 13) for both training and testing trials types. This large longitudinal sample size (3,240 trials total per participant) was chosen to focus our analyses on data at the level of individual participants (see *Statistical Analyses* below).

On each trial, a single grating was presented for 3s, followed by a 3s inter-trial interval. All runs began with a 10s fixation period and ended with a 1 min rest period. The training runs lasted for 5 mins and 43 secs, and the test runs lasted for 10 mins and 13s. Due to experimenter error during data acquisition, a portion of training trials were lost for participants 1 and 2. To compensate for the reduced number of training trials, we collected an additional session of data from subject 2, resulting in about 123 training trials and 112 testing trials per stimulus. For subject 1, we simply set aside half of the testing trials for training purposes and used the other half for testing; the number of testing trials for non-trained values of the context dimension remained the same as for all other participants.

The participants’ task was to look at a small black ring presented in the center of the screen [similar to 18]. The black ring had a small gap that randomly switched position throughout the trial. Participants were asked to continuously report the side of the gap (left or right) by pressing the corresponding button. The task had the purpose of forcing participants to fixate at the center of the screen, and to draw attention away from the stimuli.

### Functional Imaging

Imaging was performed with a Siemens Magnetom Prisma 3T whole-body MRI system located at the Center for Imaging Science, Florida International University. A volume RF coil (transmit) and a 32-channel receive array were used to acquire both functional and anatomical images. Each subject participated in four identical MRI sessions. During each session, a high-resolution 3D anatomical T1-weighted volume (MPRAGE; TR 2.4s; TI 1.1s; TE 2.9 ms; flip angle 7°; voxel size 1*×*1*×*1 mm; FOV 256 mm; 176 sagittal slices) was obtained, which served as the reference volume to align all functional images. During the main experiment, functional images were collected using a T2*-weighted EPI sequence (TR 1.5 s; TE 30 ms; flip angle 52°; sensitivity encoding with acceleration factor of 4). We collected 60 transversal slices, with resolution of 2.4*×*2.4*×*2.4 mm, and FOV of 219mm. The first six volumes in each run were discarded to allow T1 magnetization to reach steady state.

### Statistical Analyses

All data analyses, including multi-voxel decoding and tests of invariance, were performed on the individual data of each participant. In designing our experiment, we favored collection of a large amount of data per participant (3,240 trials, about 8 hours of scanning) rather than a large number of participants. Each separate analysis can be considered a replication of a single-subject experiment. With our sample sizes (*n*=135 per stimulus), our tests can detect a 6% difference from chance in classifier performance with 85% power, an 8% drop in classifier performance with >80% power, and kernel density estimate error is maximally reduced, according to simulation studies [43].

#### Region of Interest

The boundaries of V1 are commonly found using a functional localizer procedure. However, previous work has shown such boundaries can be accurately estimated from cortical folds, without the need for a functional localizer [48]. Additionally, evidence shows that the definition of V1 boundaries using the algorithm proposed by **(author?)** [48] has a precision that is equivalent to 10-25 minutes of functional mapping [49]. Therefore, we applied the **(author?)** [48] algorithm, implemented in Freesurfer 6.0 [50], to the anatomical T1-weighted images, to define the boundaries of V1 in each participant and obtain an ROI mask. The obtained V1 mask was then converted into a binary mask, and transformed to the individual’s functional scan space (the averaged volume of the first functional run was used as a target) using linear registration with FLIRT. An example mask, obtained from one participant in the study, is displayed in Figure 14.

**Figure 13:**
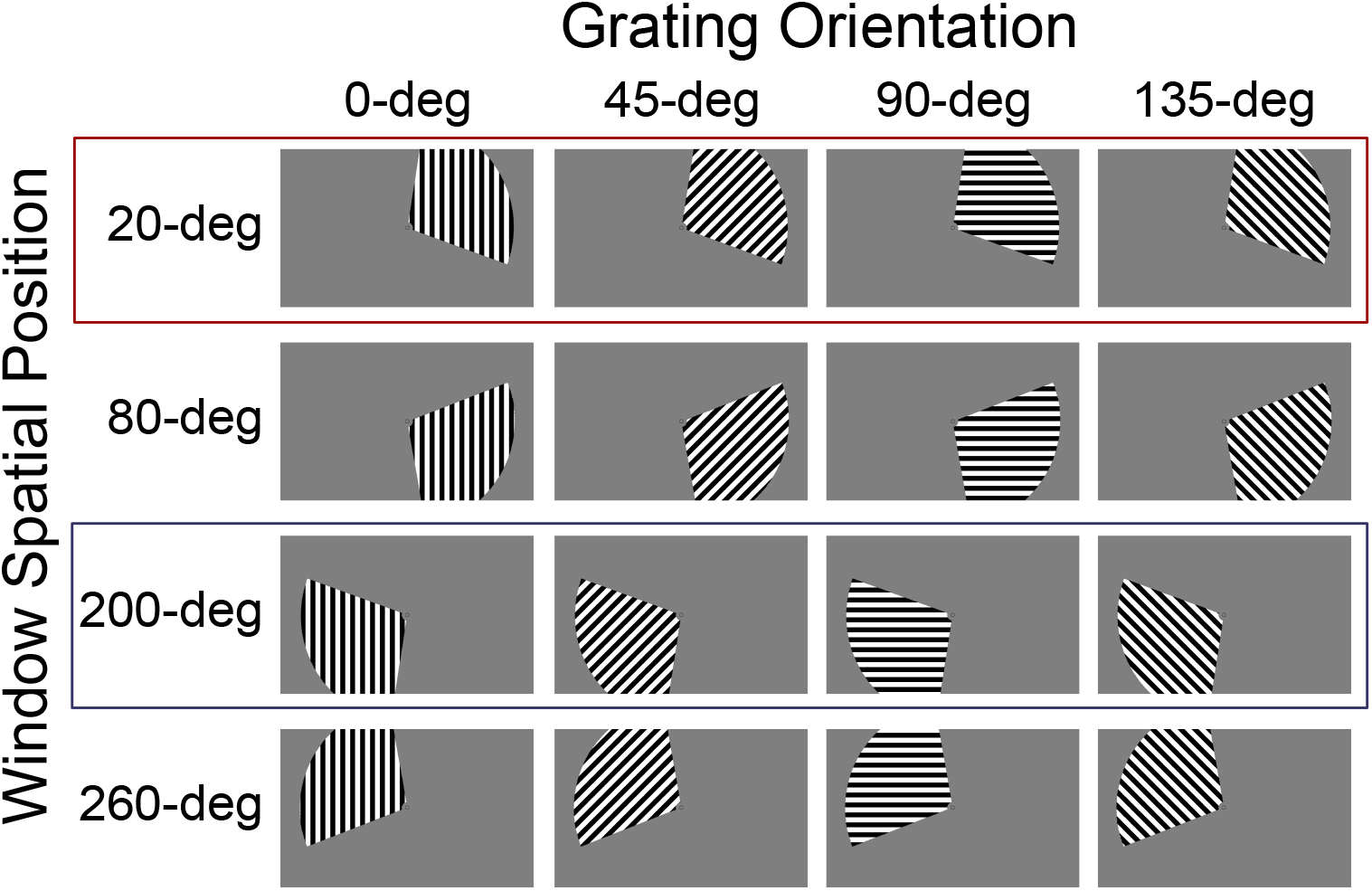
Stimuli were composed of oriented gratings (dimension 1) presented in a windowed spatial position (dimension 2). Each trial consisted of a single combination of orientated gratings and spatial position. Training runs were composed of stimuli presented only in spatial positions 20° and 200° (highlighted through red and blue boxes). Testing runs included all sixteen stimulus combinations.

**Figure 14:**
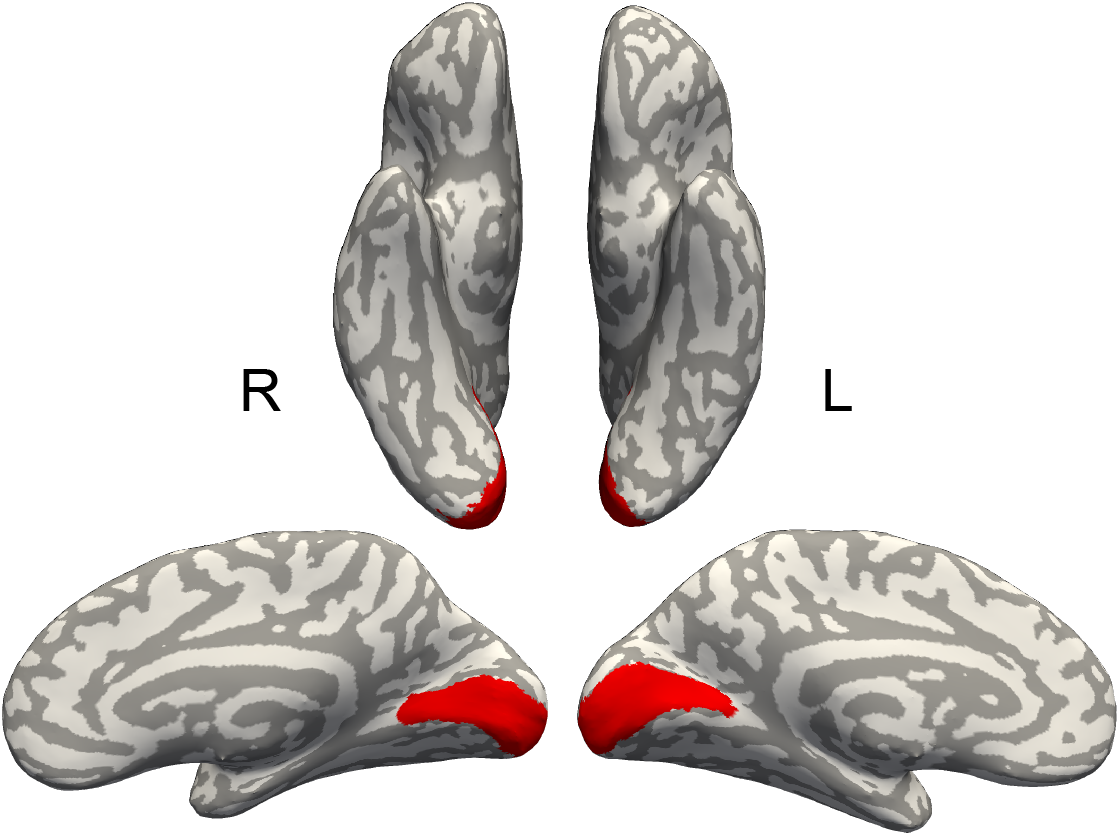
Example V1 mask obtained from one participant in this study.

#### BOLD Data Preprocessing

Data were processed and analyzed using *nipype* Python wrappers for FSL [51, 52]. Basic preprocessing of functional data included skull stripping, slice time correction, and head motion correction using MCFLIRT. All functional runs for a given subject were then aligned to an averaged volume of the first functional run for the same subject. This step ensured that the entire time-series for each subject lay in the same co-ordinate space. The aligned time-series was then concatenated into a single time-series file for further processing. The concatenated series for each subject was de-trended using a Savitzky-Golay filter with a polynomial order of 3 and a window length of 81 secs [53].

#### Deconvolution

Using the obtained V1 mask, time-series from V1 voxels were extracted for further analysis. Single-trial activity estimates were obtained via a data-driven deconvolution technique in which deconvolved neural activation values and a model of the hemodynamic response function (HRF) are estimated together [53]. Unlike other methods that hold the shape of the HRF constant across voxels, this technique allows the shape of the HRF to be different in each voxel, resulting in more accurate activity estimates. The model is implemented via the *hrf_estimation* Python package v. 1.1 (https://pypi.org/project/hrf_estimation/). The *hrf_estimation* package presents 10 different options for HRF modeling, with varying options for the HRF basis function and for the General Linear Model estimation technique. To select the optimal combination of HRF and estimation method, we performed a cross-validated decoding analysis using data from the training runs of a single participant (data from the testing runs was not used in this pre-analysis). First, we generated activity estimates from all possible model combinations (estimation method and HRF). Then, for each model, we trained and tested an SVM classifier to decode orientations from a portion of the training set, and tested the classifier with the remaining data. We chose the Rank-1 General Linear Model with a 3-basis-functions HRF model, based on the fact that it yielded the highest testing accuracy score.

#### Decoding Analysis

To decode stimulus types based on voxel-wise activity patterns, we used a Nu-support vector machine (NuSVC) classifier with a linear basis function implemented via the Python package *scikit-learn* v. 0.19.1 [54]. We used the de-convolved activity patterns from V1 voxels as inputs to the classifier, while trial-specific stimulus values (either orientation or spatial position) were provided as labels.

To decode orientation, we employed two separate classifiers, corresponding to the two different spatial positions (context dimension) at which the oriented gratings were presented during the training runs of the experiment (see Figure 13). Each classifier was trained to decode grating orientation (0°, 45°, 90°, and 135°) using only trials in which a specific spatial position was presented. However, the classifier was then tested with data collected from independent test runs at all four spatial positions. This resulted in an accuracy estimate at the training position, as well as at the other three spatial positions. For example, to train the first classifier, we gathered all trials that were presented at spatial position 20°. After normalizing the data, the classifier was trained using leave-one-run-out cross-validation with data from the training runs. Cross-validation was used to optimized the *Nu* parameter of the classifier, to obtain the highest accuracies within the training set. A new classifier was then trained on all the training data using the chosen *Nu* parameter. This classifier was then tested with data from testing runs.

To decode spatial position, we employed four separate classifiers corresponding to the four levels of grating orientation (context dimension) that were presented during the training runs of the experiment. Each classifier was trained to decode spatial position (top-right vs bottom-left, see boxed stimuli in Figure 13) using only trials in which a specific grating orientation was presented. However, the classifier was then tested with data collected from independent test runs across all levels of grating orientation. This resulted in an accuracy estimate for the training grating orientation, as well as the other three levels of grating orientation. As in the orientation decoding procedure, we divided the data into independent training and test sets, performed normalization, and optimized the classifier’s *Nu* parameter via leave-one-run-out cross-validation. One important difference is that spatial position decoding involved a two-class classification problem, where the classifiers had to discriminate between 20° and 200° spatial position of the stimulus window (the only two positions presented during training trials, see boxed stimuli in Figure 13). As the classifier was not trained to classify the 80° or 260° spatial positions, we dropped those trials from the testing data set in this analysis. This ensured that the model fitting and testing procedures remained consistent across both decoding analyses.

#### fMRI Decoding Tests

Many possible operational tests of invariance can be proposed based on the results of a decoding study, but we applied three of them to our data: the cross-classification test, the classification accuracy invariance test, and the decoding separability test. All tests where applied to the results of the two decoding analyses above: decoding of orientation and spatial position. In the descriptions below, the target dimension refers to the decoded stimulus values, and the context dimension refers to the stimulus values irrelevant for decoding that only changed from training to testing.

All the tests described below were implemented in Python expanded with *SciPy* v. 1.1.0 (https://www.scipy.org/) and *Statsmodels* v. 0.9.0 (https://www.statsmodels.org/). Plots were created using the *Matplotlib* library v. 2.2.2 (https://matplotlib.org/)

##### Cross-classification Test

The cross-classification invariance test is a well-known test in the literature that is meant to provide evidence for invariant representations directly from voxel-level activity estimates [e.g., 1, 2, 3]. As shown in Figure 2, the first step in this test is to train a classifier or another procedure to decode a particular stimulus feature from patterns of fMRI activity observed across voxels. In the figure, the feature that is being decoded is whether a face presented from a frontal viewpoint is male or female. The second step is to test the same classifier with new patterns of fMRI activity, this time obtained from presentation of the same stimuli changed in a context dimension, such as head orientation. If accuracy with this test data is higher than chance, then the conclusion is that that an invariant neural representation of the target feature is stored in the area from which the fMRI activity was obtained.

We implement the cross-classification invariance test by training a linear SVM, as described above, to classify levels of the target dimension (grating orientation or spatial position) while holding the level of the context dimension constant. For example, to decode orientation we start by training the SVM classifier to predict orientation in a given spatial position. Then, we test the accuracy of the classifier with data from independent test sets at the training position, as well as three other spatial positions (i.e., different levels of the context dimension). We tested whether each of these accuracies was above the chance level of 25% correct using a binomial test, and corrected the resulting *p*-values for multiple comparisons using the Holm-Sidak method. If classification accuracy was above chance at any of the testing levels of the context dimension, then the cross-classification invariance test concludes that there is evidence for invariance of the target dimension from the context dimension.

##### Classification Accuracy Invariance Test

Classification accuracy invariance is defined as the case where the probability of correct classification is exactly the same across all levels of the context dimension. This test is theory-driven, as it was suggested by neurocomputational theory as a valid way to obtain information about invariance [4]. Still, we are aware of at least one prior study using a version of this test to obtain evidence of position-dependent encoding of object category information in lateral occipital cortex, and of position-dependent encoding of face viewpoint information in right fusiform face area [6].

This test uses the same estimates of classification accuracy described for the cross-classification test, but uses them to check whether there is a significant drop in performance from the training to the testing context values. We first performed an omnibus Chi-Square test of the null hypothesis that accuracy does not depend on level of the context dimension. In addition, we tested accuracy at each testing context value against the training context value using a pairwise *z* test for proportions, and corrected the resulting *p*-values for multiple comparisons using the Holm-Sidak method.

##### Decoding Separability Test

Linear classifiers–like the linear SVM used here–perform classification of a new data point by computing a decision variable *z*, representing the distance of the data point from the classifier’s hyperplane separating two classes. When the decision variable is larger than some criterion value (usually zero), the output is one class, whereas when the decision variable is smaller than the criterion the output is the other class.

This suggests that a better test of invariance should use not only classification accuracy scores, but rather the full distribution of such decision variables, or *decoding distribution*. Decoding separability is defined as the case where the decoding distribution of a stimulus does not change across different levels of the context dimension (Figure 15). In Figure 15, the decoding distribution for a female face is shown for different viewpoints (context dimension). In this case, decoding separability is said to fail when the estimated decoding distribution for a target stimulus feature (e.g., maleness) is significantly different across any two levels of the context dimension (viewpoint). The distance between the two distributions of interest provides a numerical estimate of deviations from decoding separability. One way to measure this distance is through the *L*1 norm:

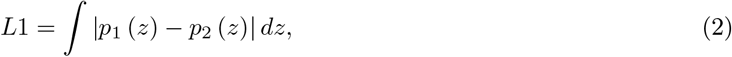

where *p*_1_ and *p*_2_ represent the distributions of decoded values at levels 1 and 2 of the context dimension, respectively. The *L*1 distance between two distributions is represented in Figure 15 by the area highlighted in light yellow.

**Figure 15:**
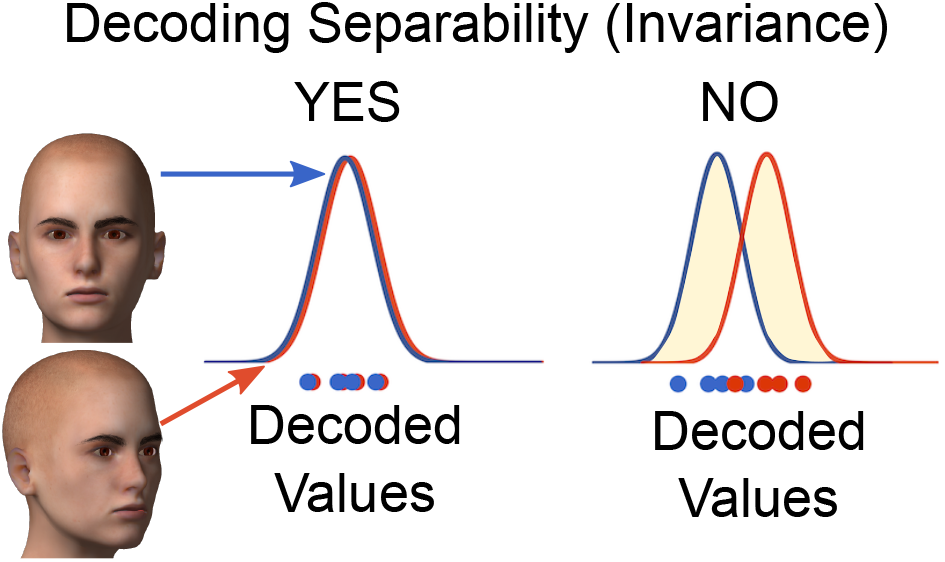
The decoding separability test. This is a theoretically-driven test of invariance that relies directly on the decoded stimulus distributions. Neural noise and random error in the decoding of stimulus values gives rise to decoding probability distributions. In this example, decoding distributions are shown for some facial identity whose viewpoint changes. If the two decoding distributions corresponding to different viewpoints are the same (left), then decoding separability is said to hold. If the decoding distributions are shifted for two levels of viewpoint (right), then decoding separability is said to fail.

For each combination of values of the relevant and context dimensions, we obtained decision variables from the trained SVM linear classifier. These decision variables were used to estimate the decoding distribution using kernel density estimates (KDEs). A gaussian kernel and automatic bandwidth determination were used as implemented in the *SciPy* function *gaussian_kde*. Let *p̂*_*ij*_ (*z*) represent the KDE for a stimulus with value *i* on the target dimension and value *j* on the context dimension, evaluated at point *z*. Each *p̂*_*ij*_ (*z*) was evaluated at values of *z* going from -3 to 6, in 0.01 steps, indexed by *k*, which were confirmed to cover the range of observed decision variable values. Then an estimate of the summed *L*1 distances indicating deviations from decoding separability was computed from all four KDEs obtained, according to the following equation:

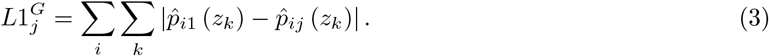

where *j* = 1 is the training level of the context dimension. The 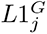 (with *G* standing for *global*) simply takes an estimate of the *L*1 distance (obtained by discretizing the continuous decision variable *z*) defined in Equation 2 for each value of the relevant dimension, and then sums them together. We computed 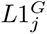 separately for each value of the context dimension, or *j* ≠ 1.

We used a permutation test to test whether each 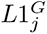 statistic was significantly larger than expected by chance. In this test, the level of the context dimension *j* was randomly re-assigned to all data points, KDEs were estimated, and the 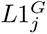 was computed according to Equation 3. This process was repeated 5,000 times, to obtain an empirical distribution for the statistic, from which accurate *p*-values were computed using the procedure proposed by [55]. The resulting *p*-values were corrected for multiple comparisons using the Holm-Sidak method.

### Simulations

The simulations described below were implemented in Python 2.6 extended with *Numpy* v. 1.16.2 (https://numpy.org/). The decoding analysis of simulated data was performed exactly as described for fMRI data in the sections *Decoding Analysis* and fMRI Decoding Tests above, with the exception that the *Nu* parameter of the SVM was set to the default value of 0.5 rather than optimized based on cross-validation.

#### Model

In our simulations, we used a standard population encoding model and a linear measurement model. Both are common choices in the computational neuroimaging literature (for a review, see [56]), both in recent simulation work [e.g., 57, 44, 25], as well as in model-based data analysis [e.g., 22, 58, 24, 41]. We assumed a circular dimension with values ranging from -90 to 90, as is the case of grating orientation, but our conclusions apply to non-circular dimensions as well.

##### Encoding Model

We used standard encoding models to represent the activity patterns of populations of neurons within a given voxel. Our encoding model was composed of several independent channels, representing any number of neurons that have similar stimulus preferences. Each channel is highly tuned to a specific value along the target stimulus dimension, such that the channel’s response becomes attenuated as we move away from the preferred value. The tuning function of a single channel is represented by a Gaussian function:

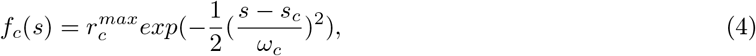

where *r^max^* represents the maximum neural activity for channel *c*, the mean *s_c_* represents the channel’s preferred stimulus, and the standard deviation *ω_c_* represents the width of the tuning function. The height of the tuning functions at any value along the stimulus dimension (i.e., *f_c_*(*s*)) represents the average response of the channels to that particular stimuli.

We assume that the response of each channel *r_c_* is a random variable with Poisson distribution:

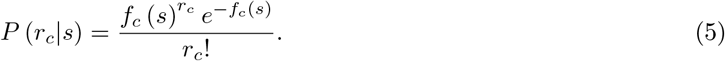

The full encoding model was composed of ten channels with activity described by Equations 4 and 5. Unless indicated otherwise below, we used a homogeneous population model, in which the parameters *s_c_* were evenly distributed across all possible values of the dimension (i.e., from -90 to 90 degrees), and other parameters were fixed to the same values for all channels: 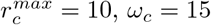.

Figure 16 shows an example of the encoding process. When a face with a value of 75% maleness is presented to the model, the channel encoding distribution produces a vector of responses. Each element in this vector corresponds to the response of a particular channel. The channels with the strongest preference for the value 75% show the highest response in this vector. Since the response of neural populations are known to be noisy, channel noise is added to each element of the response vector. The final output is a noisy vector of channel responses that change slightly for repeated presentations of the same stimulus.

**Figure 16:**
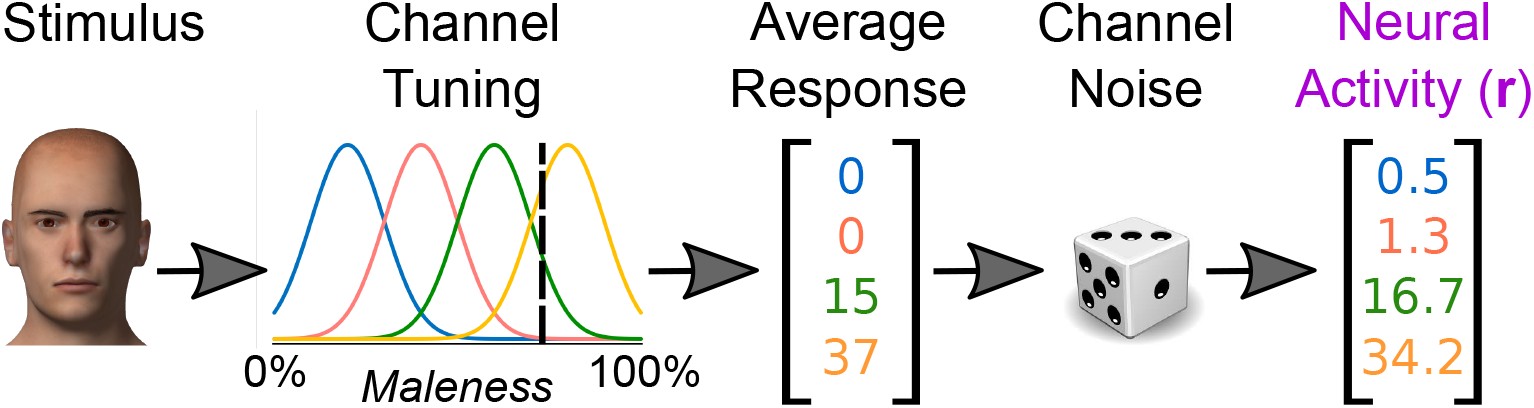
Population encoding model. This figures illustrates how stimulus information can be encoded by a set of neurons in the brain. The encoding model consists of a set of channels that are tuned to specific stimulus values along a given dimension (e.g., maleness). When a stimulus with a particular value on the maleness dimension is presented, the channels respond according to their stimulus preferences. The channel responses are then perturbed by random channel noise. The final output represents a vector of noisy firing rates in response to a particular stimulus.

##### Measurement Model

Because neuroimaging studies produce only indirect measures of neural activity, a measurement model is required to link the neural responses of the encoding model with voxel-wise activity values. The measurement model is described by the following equation:

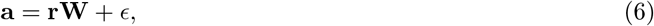

where **a** is a row vector of voxel activity values, **r** is a row vector of neural responses sampled from the encoding model (i.e., from Equation 5), **W** is a weight matrix were each column **w***_v_* represents the linear measurement model for a different voxel *a_v_*, and *E* is a random normal row vector with mean **0** and covariance matrix with *σ* in the diagonal and zeros elsewhere. The value of *σ* was varied in Simulation 1 and was fixed to 5 in Simulation 2 (see below).

Equation 6 indicates that the activity in each voxel is a linear combination of neural channel responses, plus some random measurement noise. As shown in Figure 17, the model for each voxel was composed of a finite number of encoding channels that independently contributed to the aggregate signal of the voxel according to a set of weights. The values of the weights were randomly and uniformly sampled from 0 to 1, and then normalized by column, so that weights in **w***_v_* would add up to one. This way, the weights can be interpreted as the relative contribution of each channel to a voxel’s activity.

**Figure 17:**
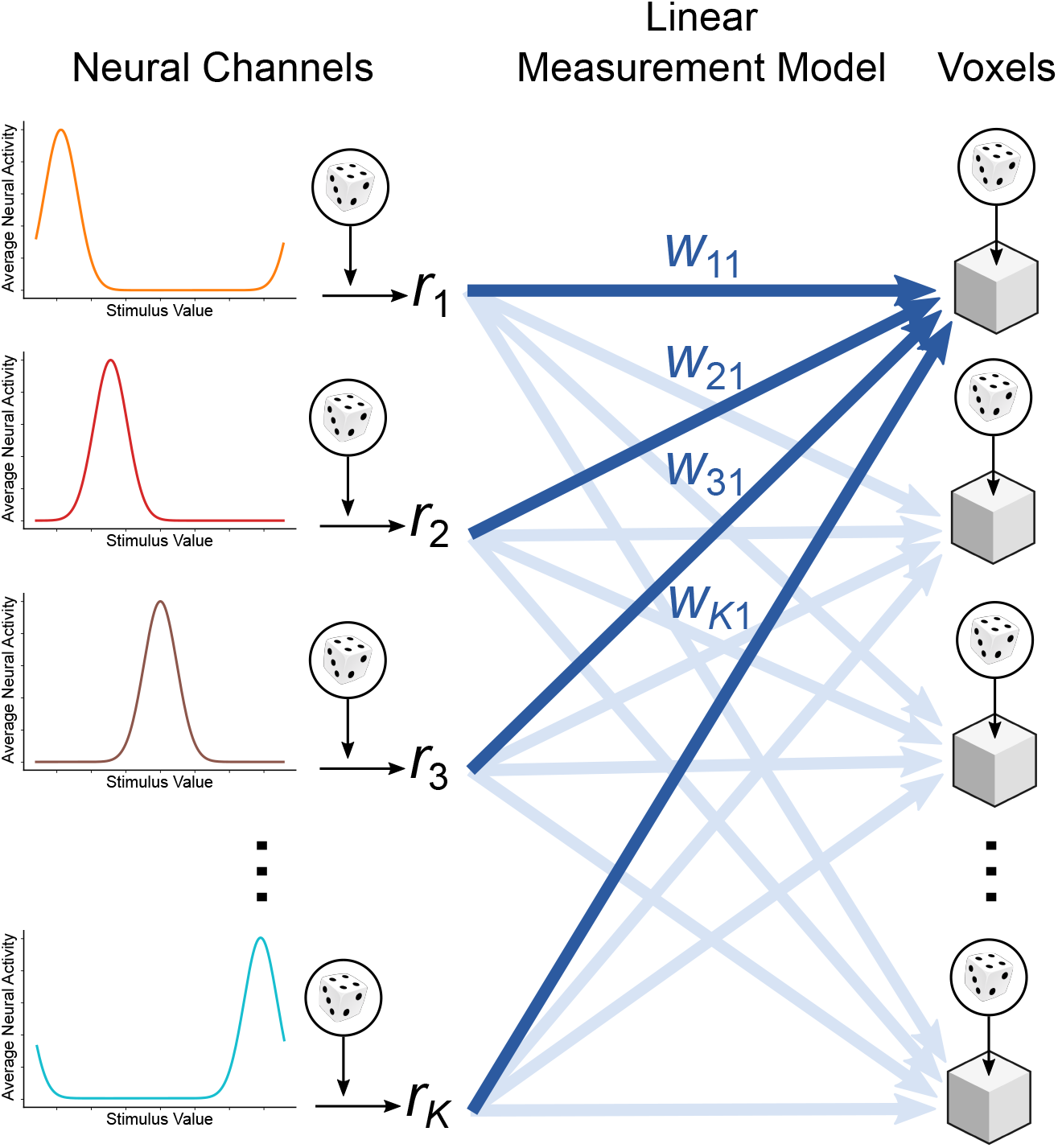
Linear measurement model. The measurement model provides a link between neural encoding channels and voxel-wise activity measures. Activity in each voxel (represented by cubes) is a linear combination of neural channel responses (**r** = [*r*_1_*, r*_2_*, r*_3_*, … r_K_*]), plus independent random measurement noise (represented by the dice next to each cube)

We simulated a total of 100 voxels. In each simulated trial, the encoding model was presented with a given stimulus and produced a random vector of neural responses **r** as explained in the previous section, which were then used as input to the measurement model to obtain a random vector of voxel activities **a**.

#### Simulation 1: False Positive Invariance Resulting From Features Of The Measurement Model

In theory, even a completely context-specific code could produce false conclusions of invariance in neuroimaging decoding studies, due to the transformation and mixing of neural responses from different populations that occurs at each voxel (see Figure 3). To provide evidence for such a general claim, we study a case of complete context-specificity in which it cannot be claimed that any amount of tolerance exists in the neural representations. The goal of the first simulation was to show that the cross-classification test can lead to a conclusion of invariance when neural representations do not satisfy any sensible definition of invariance or tolerance. As shown in Figure 18, the model underlying the simulation was created so that the encoding of the target dimension (e.g., orientation) was completely different across levels of the context dimension (e.g., spatial position). That is, two separate encoding models were created to represents the levels of the context dimension. The first level model consisted of a homogeneous population encoding model. To make sure that there was no invariance across levels of the context dimension, the encoding model for the second level of the context dimension was composed of channels whose tuning parameters were completely randomized. For each channel, the position parameter *s_c_* was randomly sampled from a uniform distribution covering all values in the dimension, *r^max^* was similarly sampled from values between 5 and 20, and *ω_c_* from values between 5 and 25. The result was a completely randomized encoding model for the second level of the context dimension, which was extremely unlikely to share any properties with the encoding model for the first level of the context dimension (compare the top and bottom encoding models in Figure 19).

**Figure 18:**
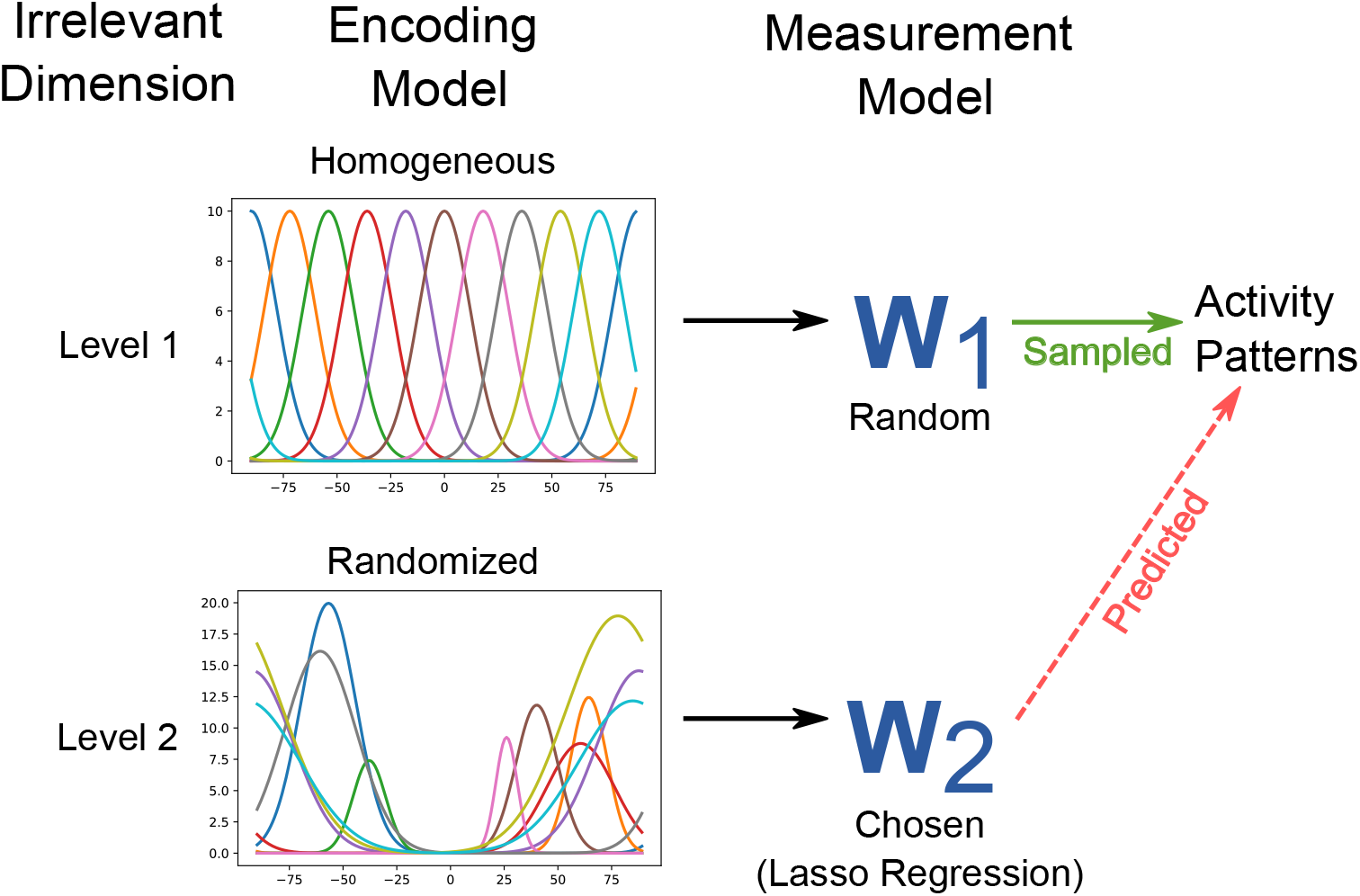
Encoding models for simulation 1. Two separate encoding models were created to represent two levels of the context dimension. The first level model (top row) consisted of homogeneously tuned encoding channels. The measurement weights for this level were selected randomly. The second level model (bottom row) consisted of randomly tuned encoding channels, ensuring a complete lack of invariance between the first and second level models. The measurement weights for this level were then chosen via lasso regression to produce similar activity patterns to that of the first level model.

As shown in the middle part of Figure 18, the measurement weights of the first level model, **W**_1_, were randomly sampled to generate activity patterns in each voxel (as explained above). On the other hand, the measurement weights for the second level model, **W**_2_, were chosen so that the activity patterns generated by any stimulus presented to this second level model would be as similar as possible as those presented to the first level model. To do this, we presented the level 1 model with the preferred stimulus of each channel *s_c_* 20 times, and each time sampled data from 100 voxels. We then presented the level 2 model with the same stimuli a single time, and recorded a vector of average responses from the encoding model using Equation 4 (i.e., neural channel responses without any noise). Finally, for each voxel, the vectors of weights in **W**_2_ were obtained via Lasso regression, where voxel-wise activity patterns produced by the first level model were used as outputs to be predicted from the average neural activities obtained from the second level encoding model. Using Lasso regression, as implemented in *sklearn*, allowed us to constrain the weights to be positive. The regularization parameter of the regression model was not optimized, but fixed to a value of 0.01.

This procedure should result in a model in which no sensible definition of invariance or tolerance holds at the level of neural responses, but that nonetheless should show some level of tolerance at the level of indirect voxel activity measures. As shown in Figure 19, each simulation started by creating such a model (step 1), and continued by sampling data from it (step 2). To get that data, we presented the model with four stimuli, with values of -45°, 0°, 45°, and 90°, and sampled voxel activity patterns from it. Each stimulus presentation was repeated 20 times. We sampled data this way both from the first and second level models constructed as indicated above. Data was sampled twice from the first level model, to obtain training and testing data sets, and only once from the second level model, to obtain a testing data set only. We then performed a cross-classification test on the resulting data (steps 3 and 4 in Figure 19), following the same procedures as with the experimental data explained above, with the exception that the *Nu* parameter of the SVM was fixed to the default value of 0.5. Each simulation was repeated 200 times. The results presented represent average statistics across all simulations, obtained from the testing data sets.

**Figure 19:**
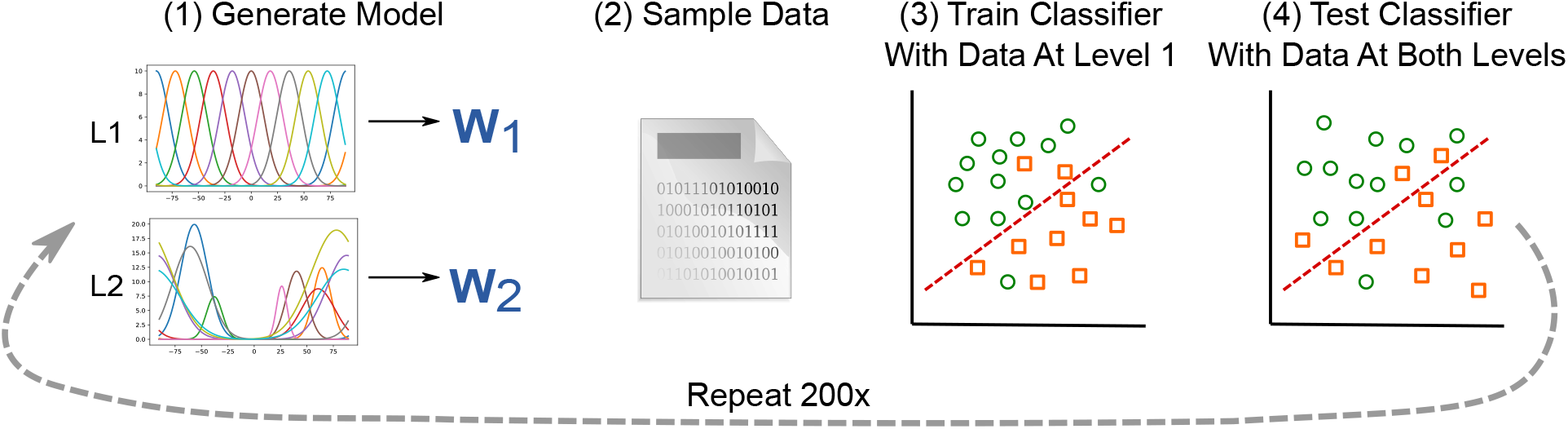
Steps taken in each repetition of simulation 1. Data was sampled from both the first and second level models. A linear SVM classifier was fit to training data from the level 1 model. Then, the fitted model was tested with independent testing data from both the first and second level models.

Finally, we repeated the group of simulations a total of 20 times, each time with a different value for the level of voxel measurement noise *σ*, going from 1 to 20.

#### Simulation 2: False Positive Invariance Resulting From Similarly Tuned Neural Subpopulations Across Contexts

Only a small proportion of all possible measurement models might be truly at play in neuroimaging studies, and those could be contained within the space of models for which false positive invariance is not an issue. Thus, we would like to strengthen our conclusions by studying a realistic encoding scenario, likely to be implemented in the brain.

There are many known cases in which neurons that are sensitive to a particular stimulus feature are spatially clustered at sub-millimeter scales. In those cases, while there is spatially distributed information about stimulus features, this information is not immediately visible at the typical resolution of an fMRI study. In cases such as these, across voxels we would expect to find relatively homogeneous distributions of selectivities. Our ability to use voxel-level decoding to detect whether and how features are encoded depends critically on small random variations in mixing; that is, in the proportion of each type of neuron present within each voxel. This sub-voxel distribution of information, which may underlie the success of many fMRI decoding studies, can also easily lead to false-positive invariance when the cross-classification test (or other tests of the null of specificity) is used in isolation. Small differences in mixing might be enough to promote above-chance decoding of a stimulus feature, because decoding algorithms are specifically trained to detect differences in the target feature. On the other hand, decoding algorithms are not trained to detect changes in context. Any small differences in mixing that might provide information about context-specificity would be lost, and the decoding algorithm would be very likely to generalize performance across changes in stimulus context.

In the present simulation, we wanted to study the sensitivity of different fMRI decoding tests to changes in mixing indicative of context-sensitivity. With this goal in mind, we created a model in which a target dimension is encoded in a completely context-specific manner, with one subpopulation of neurons responding whenever the context dimension is at level 1, and a different subpopulation of neurons responding whenever the context dimension is at level 2. However, both subpopulations encoded the target dimension in a similar way. To do this, the measurement model of each voxel was obtained as shown in Figure 20. The weights **w***_v_* for level 1 of the context dimension were randomly generated, as explained above (step 1 in Figure 20). To create the measurement model for level 2 of the context dimension, we first obtained a vector **e** of random values sampled from a normal distribution with mean zero and standard deviation equal to *σ***_e_**. This random vector was added to **w***_v_* (step 2), and then the values were made positive through rectification and normalized to add up to one (step 3), producing the final set of weights for level 2 of the context dimension. Once the model was generated, the simulation was carried out following the same other steps as in Simulation 1, numbered 2 to 4 in Figure 19. The only difference was that *σ* = 5 in the measurement model, whereas the value of *σ***_e_** was varied from 0 to 0.5. Note that this implies that, at the highest values of *σ***_e_**, the standard deviation of the changes in weights in the measurement model was 500% the average value of those weights (0.1). This ensured that in the final models the contribution of each neuron type (e.g., neurons selective to a value of 0 in the target dimension) was widely different across levels of the context dimension.

**Figure 20:**
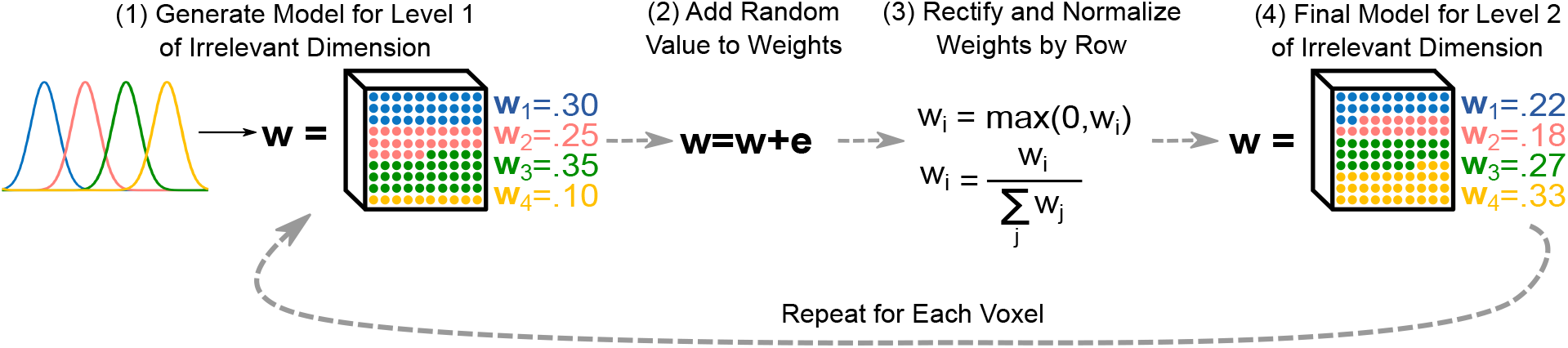
Encoding model for simulation 2. The encoding model for the first level was generated in the same manner as in simulation 1 (see Figure 18). For the second level model, the channels weights within each voxel were perturbed by a vector of random values sampled from a normal distribution, and were then rectified and normalized to make sure that they were positive and added up to one for each voxel.

## Acknowledgements

This work was supported in part by the National Science Foundation under grant No. 2020982 to Fabian A. Soto. Any opinions, findings, and conclusions or recommendations expressed in this material are those of the author(s) and do not necessarily reflect the views of the National Science Foundation. We thank Jason Hays for developing and teaching us how to use the Python package used for our simulations (PEMGUIN). *S.D.G*.

