## Supplementary Material for "Improving the validity of neuroimaging decoding tests of invariant and configural neural representation"

| | Sub #1 | | Sub #2 | | Sub #3 | | Sub#4 | | Sub#5 | |
|---|---|---|---|---|---|---|---|---|---|---|
| | Accuracy | P-value | Accuracy | P-value | Accuracy | P-value | Accuracy | P-value | Accuracy | P-value |
| ***Spatial Position Decoding*** | | | | | | | | | | |
| *Training Orientation 0-deg* | | | | | | | | | | |
| Testing Orientation 0-deg | 95.59 | <.001 | 100.00 | <.001 | 95.19 | <.001 | 81.04 | <.001 | 95.19 | <.001 |
| Testing Orientation 45-deg | 99.26 | <.001 | 100.00 | <.001 | 93.36 | <.001 | 82.53 | <.001 | 94.09 | <.001 |
| Testing Orientation 90-deg | 97.76 | <.001 | 100.00 | <.001 | 95.93 | <.001 | 81.18 | <.001 | 93.70 | <.001 |
| Testing Orientation 135-deg | 95.52 | <.001 | 100.00 | <.001 | 93.68 | <.001 | 84.87 | <.001 | 96.28 | <.001 |
| *Training Orientation 45-deg* | | | | | | | | | | |
| Testing Orientation 45-deg | 100.00 | <.001 | 100.00 | <.001 | 95.94 | <.001 | 76.21 | <.001 | 94.09 | <.001 |
| Testing Orientation 0-deg | 96.32 | <.001 | 100.00 | <.001 | 94.81 | <.001 | 75.46 | <.001 | 96.29 | <.001 |
| Testing Orientation 90-deg | 97.76 | <.001 | 100.00 | <.001 | 96.29 | <.001 | 81.92 | <.001 | 93.70 | <.001 |
| Testing Orientation 135-deg | 96.27 | <.001 | 100.00 | <.001 | 94.05 | <.001 | 76.38 | <.001 | 95.91 | <.001 |
| *Training Orientation 90-deg* | | | | | | | | | | |
| Testing Orientation 90-deg | 98.51 | <.001 | 100.00 | <.001 | 94.07 | <.001 | 80.07 | <.001 | 94.07 | <.001 |
| Testing Orientation 45-deg | 99.26 | <.001 | 100.00 | <.001 | 94.46 | <.001 | 79.18 | <.001 | 95.57 | <.001 |
| Testing Orientation 135-deg | 94.78 | <.001 | 100.00 | <.001 | 93.68 | <.001 | 80.44 | <.001 | 96.28 | <.001 |
| Testing Orientation 0-deg | 95.59 | <.001 | 100.00 | <.001 | 96.96 | <.001 | 79.55 | <.001 | 95.19 | <.001 |
| *Training Orientation 135-deg* | | | | | | | | | | |
| Testing Orientation 135-deg | 96.27 | <.001 | 100.00 | <.001 | 94.42 | <.001 | 83.76 | <.001 | 96.28 | <.001 |
| Testing Orientation 0-deg | 95.59 | <.001 | 100.00 | <.001 | 96.29 | <.001 | 83.27 | <.001 | 95.56 | <.001 |
| Testing Orientation 90-deg | 97.76 | <.001 | 100.00 | <.001 | 95.93 | <.001 | 84.13 | <.001 | 95.56 | <.001 |
| Testing Orientation 45-deg | 99.26 | <.001 | 100.00 | <.001 | 95.20 | <.001 | 80.29 | <.001 | 94.46 | <.001 |
| ***Orientation Decoding*** | | | | | | | | | | |
| *Training Window 20-deg* | | | | | | | | | | |
| Testing Window 20-deg | 24.81 | .959 | 40.89 | <.001 | 38.70 | <.001 | 28.62 | .265 | 34.07 | <.001 |
| Testing Window 80-deg | 23.70 | .988 | 16.00 | .999 | 30.37 | .008 | 23.79 | .698 | 26.11 | .643 |
| Testing Window 260-deg | 20.55 | .999 | 27.11 | .414 | 28.52 | .068 | 30.48 | .093 | 22.96 | .970 |
| Testing Window 200-deg | 15.56 | .999 | 18.89 | .999 | 27.22 | .127 | 25.65 | .670 | 23.33 | .970 |
| *Training Window 200-deg* | | | | | | | | | | |
| Testing Window 200-deg | 35.19 | <.001 | 54.44 | <.001 | 35.74 | <.001 | 32.71 | .011 | 34.07 | <.001 |
| Testing Window 260-deg | 27.22 | .335 | 20.44 | .999 | 28.52 | .099 | 24.91 | .786 | 25.92 | .544 |
| Testing Window 80-deg | 22.22 | .996 | 30.22 | .021 | 26.11 | .290 | 28.25 | .326 | 24.81 | .556 |
| Testing Window 20-deg | 18.52 | .999 | 19.11 | .999 | 27.41 | .204 | 23.05 | .789 | 27.96 | .177 |

Table 1: Detailed results of the cross-classification test. P-values have been corrected for multiple comparisons using the Holm-Sidak method.

| | Sub #1 | | Sub #2 | | Sub #3 | | Sub#4 | | Sub#5 | |
|---|---|---|---|---|---|---|---|---|---|---|
| | Statistic | P-value | Statistic | P-value | Statistic | P-value | Statistic | P-value | Statistic | P-value |
| ***Spatial Position Decoding*** | | | | | | | | | | |
| *Training Orientation 0-deg* | | | | | | | | | | |
| Omnibus Test | $\chi^2(3)$= 4.65 | .19 | - | - | $\chi^2(3)$= 2.34 | .505 | $\chi^2(3)$= 1.76 | .623 | $\chi^2(3)$= 2.22 | .529 |
| 0-deg vs. 45-deg | $z$= -1.91 | .16 | - | - | $z$= .91 | .739 | $z$= -.45 | .881 | $z$= .56 | .836 |
| 0-deg vs. 90-deg | $z$= .03 | .98 | - | - | $z$= -.42 | .739 | $z$= -1.18 | .555 | $z$= .75 | .836 |
| 0-deg vs. 135-deg | $z$= -.99 | .54 | - | - | $z$= .76 | .739 | $z$= -.04 | .967 | $z$= -.63 | .836 |
| *Training Orientation 45-deg* | | | | | | | | | | |
| Omnibus Test | $\chi^2(3)$= 5.30 | .15 | - | - | $\chi^2(3)$= 1.91 | .590 | $\chi^2(3)$= 4.12 | .248 | $\chi^2(3)$= 2.85 | .416 |
| 45-deg vs. 0-deg | $z$= 2.26 | .07 | - | - | $z$= .62 | .782 | $z$= 20 | .975 | $z$= -1.19 | .546 |
| 45-deg vs. 90-deg | $z$= 1.75 | .08 | - | - | $z$= -.21 | .831 | $z$= -1.63 | .278 | $z$= .19 | .849 |
| 45-deg vs. 135-deg | $z$= 2.27 | .07 | - | - | $z$= 1.01 | .677 | $z$= -.048 | .975 | $z$= -.97 | .555 |
| *Training Orientation 90-deg* | | | | | | | | | | |
| Omnibus Test | $\chi^2(3)$= 6.73 | .08 | - | - | $\chi^2(3)$= 2.13 | .546 | $\chi^2(3)$= .16 | .984 | $\chi^2(3)$= 1.53 | .675 |
| 90-deg vs. 45-deg | $z$= -.59 | .55 | - | - | $z$= -.19 | .976 | $z$= .26 | .992 | $z$= -.79 | .677 |
| 90-deg vs. 135-deg | $z$= 1.69 | .25 | - | - | $z$= .19 | .976 | $z$= -.11 | .992 | $z$= -1.19 | .546 |
| 90-deg vs. 0-deg | $z$= 1.41 | .29 | - | - | $z$= -1.21 | .539 | $z$= .15 | .992 | $z$= -.57 | .677 |
| *Training Orientation 135-deg* | | | | | | | | | | |
| Omnibus Test | $\chi^2(3)$= 4.04 | .26 | - | - | $\chi^2(3)$= 1.28 | .734 | $\chi^2(3)$= 1.74 | .628 | $\chi^2(3)$= 1.05 | .789 |
| 135-deg vs. 0-deg | $z$= .28 | .78 | - | - | $z$= -1.03 | .659 | $z$= .15 | .985 | $z$= .43 | .891 |
| 135-deg vs. 90-deg | $z$= -.72 | .72 | - | - | $z$= -.81 | .659 | $z$= -.11 | .985 | $z$= .43 | .891 |
| 135-deg vs. 45-deg | $z$= 1.67 | .26 | - | - | $z$= -.41 | .683 | $z$= 1.05 | .648 | $z$= 1.01 | .678 |
| ***Orientation Decoding*** | | | | | | | | | | |
| *Training Window 20-deg* | | | | | | | | | | |
| Omnibus Test | $\chi^2(3)$= 14.49 | .002 | $\chi^2(3)$= 87.89 | <.001 | $\chi^2(3)$= 20.12 | <.001 | $\chi^2(3)$= 3.65 | .302 | $\chi^2(3)$= 22.11 | <.001 |
| 20-deg vs. 80-deg | $z$= .35 | .727 | $z$= 8.28 | <.001 | $z$= 2.88 | .004 | $z$= 1.27 | .493 | $z$= 2.85 | .004 |
| 20-deg vs. 260-deg | $z$= 1.38 | .307 | $z$= 4.36 | <.001 | $z$= 3.54 | <.001 | $z$= -.47 | .684 | $z$= 4.04 | <.001 |
| 20-deg vs. 200-deg | $z$= 3.19 | .004 | $z$= 7.21 | <.001 | $z$= 4.01 | <.001 | $z$= .78 | .684 | $z$= 3.90 | <.001 |
| *Training Window 200-deg* | | | | | | | | | | |
| Omnibus Test | $\chi^2(3)$= 30.83 | <.001 | $\chi^2(3)$= 168.77 | <.001 | $\chi^2(3)$= 14.49 | .002 | $\chi^2(3)$= 7.33 | .062 | $\chi^2(3)$= 13.65 | .003 |
| 200-deg vs. 260-deg | $z$= 2.33 | .019 | $z$= 10.54 | <.001 | $z$= 2.54 | .011 | $z$= 1.99 | .089 | $z$= 2.92 | .006 |
| 200-deg vs. 80-deg | $z$= 3.94 | <.001 | $z$= 7.35 | <.001 | $z$= 3.42 | .002 | $z$= 1.12 | .261 | $z$= 3.34 | .003 |
| 200-deg vs. 20-deg | $z$= 5.23 | <.001 | $z$= 10.99 | <.001 | $z$= 2.95 | .006 | $z$= 2.49 | .037 | $z$= 2.17 | .029 |

Table 2: Detailed results of the classification invariance test. The symbol "-" indicates that the test could not be performed due to lack of variability in the results (all accuracies at the ceiling of 100%). P-values have been corrected for multiple comparisons using the Holm-Sidak method.

| | Sub #1 | | Sub #2 | | Sub #3 | | Sub#4 | | Sub#5 | |
|---|---|---|---|---|---|---|---|---|---|---|
| | $L1_j^G$ Statistic | P-value | $L1_j^G$ Statistic | P-value | $L1_j^G$ Statistic | P-value | $L1_j^G$ Statistic | P-value | $L1_j^G$ Statistic | P-value |
| ***Spatial Position Decoding*** | | | | | | | | | | |
| *Training Orientation 0-deg* | | | | | | | | | | |
| 0-deg vs. 45-deg | 43.25 | .019 | 63.33 | <.001 | 28.73 | .054 | 31.56 | .023 | 50.24 | <.001 |
| 0-deg vs. 90-deg | 45.15 | .019 | 45.55 | <.001 | 27.37 | .061 | 28.67 | .034 | 50.44 | <.001 |
| 0-deg vs. 135-deg | 43.41 | .019 | 50.83 | <.001 | 26.36 | .061 | 20.09 | .191 | 46.74 | <.001 |
| *Training Orientation 45-deg* | | | | | | | | | | |
| 45-deg vs. 0-deg | 42.40 | .037 | 27.38 | .094 | 21.01 | .156 | 20.51 | .161 | 48.39 | <.001 |
| 45-deg vs. 90-deg | 36.48 | .052 | 49.15 | <.001 | 33.19 | .010 | 39.07 | <.001 | 40.08 | <.001 |
| 45-deg vs. 135-deg | 38.44 | .052 | 53.43 | <.001 | 29.71 | .034 | 27.24 | .056 | 37.13 | .001 |
| *Training Orientation 90-deg* | | | | | | | | | | |
| 90-deg vs. 45-deg | 45.96 | .017 | 22.69 | .219 | 32.41 | .014 | 30.42 | .033 | 48.96 | <.001 |
| 90-deg vs. 135-deg | 31.63 | .103 | 37.83 | .007 | 52.84 | <.001 | 30.63 | .033 | 54.35 | <.001 |
| 90-deg vs. 0-deg | 39.77 | .036 | 38.86 | .008 | 30.16 | .014 | 25.68 | .040 | 47.40 | <.001 |
| *Training Orientation 135-deg* | | | | | | | | | | |
| 135-deg vs. 0-deg | 50.46 | .004 | 29.94 | .040 | 26.47 | .039 | 27.17 | .082 | 54.94 | <.001 |
| 135-deg vs. 90-deg | 37.26 | .053 | 44.37 | <.001 | 49.39 | <.001 | 27.32 | .082 | 49.43 | <.001 |
| 135-deg vs. 45-deg | 38.60 | .053 | 50.29 | <.001 | 29.00 | .039 | 18.93 | .248 | 45.85 | <.001 |
| ***Orientation Decoding*** | | | | | | | | | | |
| *Training Window 20-deg* | | | | | | | | | | |
| 20-deg vs. 80-deg | 104.51 | <.001 | 101.05 | .004 | 65.32 | .208 | 74.28 | .713 | 72.73 | .163 |
| 20-deg vs. 260-deg | 76.81 | .307 | 90.32 | .016 | 90.96 | .008 | 87.72 | .504 | 49.88 | .688 |
| 20-deg vs. 200-deg | 71.37 | .199 | 187.63 | <.001 | 109.78 | <.001 | 63.73 | .719 | 86.35 | .024 |
| *Training Window 200-deg* | | | | | | | | | | |
| 200-deg vs. 260-deg | 114.29 | .006 | 281.98 | <.001 | 79.34 | .031 | 106.29 | .082 | 78.88 | .068 |
| 200-deg vs. 80-deg | 102.44 | .001 | 208.15 | <.001 | 105.77 | <.001 | 92.98 | .125 | 95.57 | .004 |
| 200-deg vs. 20-deg | 91.25 | .006 | 314.68 | <.001 | 84.35 | .026 | 106.27 | .082 | 69.58 | .120 |

Table 3: Detailed results of the decoding separability test. P-values have been corrected for multiple comparisons using the Holm-Sidak method.
